# Fitness of a clonal population can be inferred from lineage trees without knowledge of the biological details

**DOI:** 10.1101/2022.09.09.507320

**Authors:** Javier Escabi, Sahand Hormoz

## Abstract

Inferring the rate at which a clonal population grows, or its fitness, is important for many biomedical applications. For example, measuring the fitness of mutated cells in a patient with cancer may provide important information about prognosis and treatment. Similarly, measuring the fitness of new viral strains that emerge during a pandemic can inform how to plan an effective response. In previous work, the lineage trees constructed from individuals randomly sampled from the population at the final time-point have been used to infer the fitness and the times at which the mutation providing the fitness advantage arose in a diverse set of systems, such as blood cancers [1], [2] and the influenza virus [3]. However, it is not clear to what extent the inferred values depend on the exact biological details assumed in the models used for the inference. In this paper we show that coalescent statistics of lineage trees are invariant to changes in key parameters underlying the expansion, such as the distribution of the number of progenies produced by each individual and heterogeneity in the expansion rate. In addition, we show that competition between drift and selection imply that the fitness of the mutated population and when the mutation occurred can be inferred without knowledge of the mutation rate per generation even though the population size itself cannot be inferred. Lastly, we show that our results also generalize to cases where multiple competing mutations result in multiple distinct subclones with different values of fitness. Taken together, our results show that inferring fitness from lineage trees is robust to most model assumptions.

## Introduction

In many biological processes, a single individual generates identical copies of itself which in turn generate additional identical copies of themselves and so on, resulting in an exponentially growing clonal population. For example, certain types of blood cancers occur when a blood stem cell accrues a mutation that gives it a selective advantage. The mutated stem cell then divides repeatedly and gives rise to an exponentially growing clone of progenies that carry the same mutation [1], [2], [4], [5]. Similarly, infections can occur when a virus infects a host cell and generates thousands of new progenies which in turn infect other cells resulting in an exponentially growing population of virus particles [6]. In these systems and others, we would like to know when the exponential growth started and the rate at which the population grew, referred to as fitness.

One way to obtain these quantities is to directly observe the expanding population. However, in most cases direct observations are not feasible –we cannot follow individual cancer cells as they divide in each patient or track the number of virus particles in a host over time. Rather, in most cases we only have access to a subset of individuals randomly sampled from the population at the final time-point. The only information that can be obtained from a sample of individuals at the final time-point is their degree of relatedness or their lineage tree. To obtain the lineage tree, we can sequence the genomes of the sampled individuals and use the pattern of randomly occurring neutral mutations accrued over time to establish who is related to whom. Here, we will show that fitness of an exponentially growing clonal population can be inferred from the lineage tree of a subset of individuals randomly sampled at the final time-point. Importantly, we will argue that fitness can be inferred accurately without knowledge of the details of the biological process that gave rise to the tree.

The biological processes giving rise to exponentially growing clonal populations vary significantly across different species and systems. For example, cancer cells undergo binary cell divisions that produce precisely two children whereas viruses infect host cells and produce thousands of progenies. Even among different clones of the same system, for example the same type of cancer initiated by the same mutation but occurring in two distinct patients, the average time between replication events, the number of progenies produced, and the timing of death events could vary as well as the degree to which these quantities fluctuate. It might be expected that the fitness inferred from a lineage tree should sensitively depend on the details of the underlying biological process that generated the tree. Surprisingly, however, we will show that the coalescence structure of lineage trees only depends on fitness and is invariant to other parameters.

To do so, we start off with the well-known observation that the coalescent structure of the lineage tree of individuals randomly sampled from a population with fixed population size undergoing neutral dynamics is given by Kingman coalescence [7]. In Kingman coalescence, the most recent common ancestor of two randomly chosen individuals occurs *T* generations back with probability 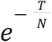, where *N* is the population size. When the population size changes across generations, this probability is given by 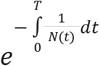. Therefore, the coalescent times of the lineage tree should allow us to infer *N*(*t*) and in turn the rate of growth of the population. For most applications, however, we cannot measure the lengths of the branches of the lineage tree in number of generations. Instead, the branch lengths are measured in units of number of mutations. Therefore, the population size can only be inferred up to a constant proportional to the mutation rate given in mutations per generation. For example, two individuals that differ by a large number of mutations can be close relatives in a small population with a high mutation rate or conversely distant relatives in a large population with a slow mutation rate.

First, we will show that fitness can be inferred without knowledge of the mutation rate per generation. This is counterintuitive because we might expect that if the population size is unknown then the rate of growth of the population should also be unknown. This is because the mutated population starts off as one individual and if its rate of growth is known then naively we should be able to determine the size of the population at any time point. As we will argue, this naive argument neglects the stochastic fluctuations in the population size caused by drift when the population size is small. With drift accounted for, fitness can be inferred without knowledge of the mutation rate.

Second, we will show that the time at which the first mutated individual appeared (time of onset) can also be inferred without knowledge of the mutation rate per generation. This is a counterintuitive result because we might expect that if the population size is not known then we cannot determine the time at which the population size was equal to one. We will show that the inferred time of onset is different from the true time on average by at most the duration of one generation. Therefore, the age of onset of the cancer mutation in a patient can be inferred accurately from the reconstructed phylogenetic tree of the cancer cells without knowing how many divisions the cancer cells have undergone.

Third, we will show that changing the distribution of the number of progenies rescales the population size (much like the mutation rate) while leaving the coalescent structure of the lineage tree invariant. Therefore, fitness can be inferred without any knowledge of the number of progenies produced at each generation and their degree of fluctuations. An expanding population of cancer cells that always generate exactly two progenies at each division will give rise to lineage trees that are identical to that of viruses that generate thousands of new progenies with large fluctuations at each generation as long as the cancer cells and the viruses have the same fitness (Figure 1). In addition, we will show that coalescent statistics of trees are invariant to fluctuations in fitness of each individual, as long as these fluctuations are independent and not passed on to the progenies. This implies that average fitness can be inferred accurately from trees even if there is heterogeneity in a population.

**Figure 1.**
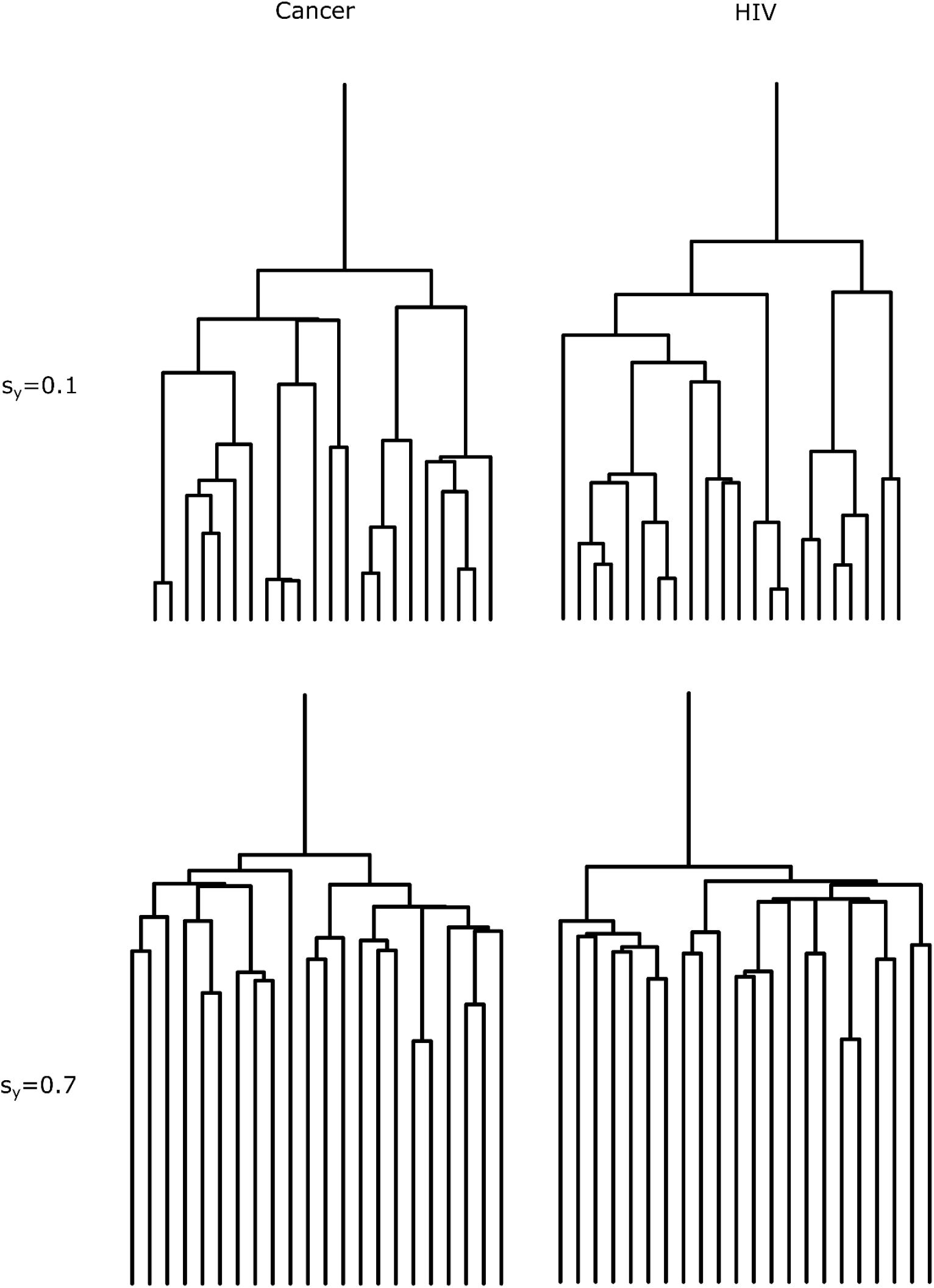
Trees constructed from simulated cancer cell divisions look the same as trees constructed from simulated replication of HIV as long as the fitness is the same. Simulated trees were constructed using realistic parameter values from MPN bone marrow cancers and HIV replication under different fitness values. 22 lineages were randomly sampled at the final time-point and the corresponding trees were plotted. The trees on the left hand side were constructed from simulated cancer expansions, while the trees on the right hand side from simulated HIV expansions. The trees on the top row have the same fitness *s*_*y*_ = 0. 1 even though the biological details differ, and have different fitness values to the trees on the bottom row which have fitness *s*_*y*_ = 0. 1. As expected the trees on the top row look different to the trees on the bottom row because their fitness values differ. However, the trees on the left column are indistinguishable from the corresponding tree on the right column because only the biological details have changed.

Lastly, we will show that these same principles apply to the lineage trees of populations with multiple competing mutations such as a cancer with multiple driver mutations or multiple competing strains of a virus. As a result, the fitness values and times of onset of each individual clone can be inferred from the lineage tree of the population without knowledge of the mutation rate, the distribution of progeny, or non-heritable fluctuations in fitness across the individuals of the population.

Taken together, our results imply that inference of fitness and time of onset from empirically observed lineage trees is robust to many details of the models used for the inference. For example, in Bayesian inference, we can simulate a specific model of an exponentially growing population for different values of fitness and retain the values that produce trees that are sufficiently close to the observed tree. Even if this model makes drastically incorrect assumptions about the biological process, for example an incorrect mutation rate, neglects heterogeneity in fitness across the population, or if it assumes that the distribution of progenies is that of a virus instead of cancer cells, the inferred fitness would still be correct.

## Description of the model

We will show that fitness and time of onset of the mutant population can be inferred correctly even when many assumptions of the model used for the inference are wrong. To do so, we will show that the coalescent statistics of the lineage tree of a subset of mutant individuals randomly selected at the final time point are invariant to changes in some of the key model parameters. We initially use the Wright-Fisher model with selection where only a single driver mutation produces the exponential growth because it is easy to change the biological assumptions of the model, extend it to more complex scenarios, and compute the coalescent statistics. In particular, the Wright-Fisher model with selection has generations that are discrete and non-overlapping which simplifies the calculations.

Our model begins with a total population of *N* identical individuals. We assume that *N* remains fixed over time. *N* can be thought of as the carrying capacity of the environment. At each generation the individuals at the current generation give birth to a new generation of *N* individuals and immediately die off. Each individual in the new generation can descend from any individual in the previous generation with equal probability. This process is iterated until at some random generation (referred to as the time of onset) one individual acquires a mutation that gives it a selective advantage *s*. From then on, the probabilities of descent change. Each individual in the new generation descends from wild-type individuals in the previous generation with probability *p*, and from mutant individuals in the previous generation with probability (1 + *s*)*p*. Note that as expected, when *s* = 0 all individuals in the previous generation are equally likely to be the parent of a given individual in the new generation. If *n* is the number of mutant individuals in the previous generation, then *p* can be derived from the condition that probabilities must sum to 1:

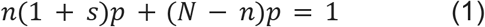

We then define *L* to be the total number of generations of the Wright-Fisher process, and *g* to be the number of generations from the time at which the mutant individual arose all the way to the *L* ^*th*^ generation at which the process terminates. Lastly, we define *a* to be the age of the population, or the amount of time (for example in years) that has elapsed from the first generation to the *L*^*th*^. Note that the number of generations per unit time is then 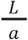.

### The intuition for why fitness can be inferred without knowledge of the number of generations

In the following section, we argue intuitively why it makes sense that we can generally infer the percent growth per year, or fitness, from the lineage tree of a random sample at the final time-point without knowing the total number of generations spanned by the lineage tree.

To begin, let’s consider a population that evolves according to the Wright-Fisher process with selection as defined in the previous section. Let’s assume that we know the total number of generations *L* and the age of the population *a*. We assume that the clone has not gone extinct so that individuals from the mutant clone can be sampled at the final time-point. We also assume that mutations are accrued at a constant rate and that each site in the genome is mutated only once. The lineage tree can then be reconstructed from the number of mutations that are shared across different individuals in the sample, for example using the UPGMA algorithm [8]. We can estimate the number of mutations per unit time as the average number of mutations in the genomes of each individual in the sample divided by a. This allows us to convert the lengths of branches of the tree from number mutations to the amount of time. Since there are fluctuations in the number of mutations accrued per individual per generation, this tree is only an estimate of the actual genealogy of the sample. Although in practice we will only be able to estimate the genealogy, we will ignore the uncertainty introduced by fluctuations in the number of mutations and assume that we have perfect knowledge of the genealogy of the sample with branch lengths measured in units of time. If *L* is known, we can use the number of generations per unit time 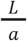 to convert the branch lengths of the tree from units of time to number of generations.

We can then ask how the structure of the reconstructed trees change when *s*, or the selective advantage, is changed for fixed *L*. A clone with selective advantage *s* grows exponentially as ∼(1 + *s*) ^*t*^ [1] where *t* is the number of generations since the mutation arose. When *s* is increased, the number of mutant individuals over time also increases, especially for larger values of t. Conversely, when *s* is decreased, the number of mutant individuals over time decreases. When the number of mutant individuals is large, the probability that the sampled individuals are closely related is small, therefore, the lineages take a longer time to coalesce (merge together) when going from the bottom of the tree to the top [7]. Conversely, when the number of mutant individuals is small, the probability that the sampled individuals are closely related is large, and so the lineages will tend to coalesce closer to the bottom of the tree. Thus different values of *s* give rise to different coalescent statistics of trees, in particular, larger *s* means that fewer coalescence events occur towards the bottom of the tree. Since we can distinguish trees with different values of *s*, in general, we can infer *s*, from observed trees whenever *L* is known.

When *L* is unknown, we cannot infer the population size because we cannot convert the lengths of branches of the tree from units of time to number of generations. We also cannot infer the selective advantage *s*, or equivalently the percent growth rate per generation, but we can still infer the percent growth rate per unit time, referred to as fitness. The distinction between these two quantities is important. For example, blood cancer cells may undergo multiple cell divisions per year on average so that the number of cells more than double in a year even though precisely two cells are produced from each cancer cell at each generation.

We now derive the relationship between *s* and fitness. If the average growth per generation of a clone is a factor 1 + *s*, and the Wright-Fisher process has gone on for *L* generations or *a* units of time, then the growth per unit time of the clone is a factor of 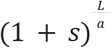, and so the fitness *s* is:

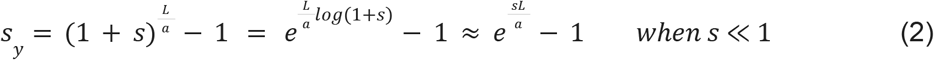

The clone then grows as ∼(1 + *s* _*y*_)^τ^, where τ is given in units of time.

Our central claim is that *s* _*y*_ can be inferred without any knowledge of *L*. To motivate this claim, we consider the average growth of the mutant clone. First, note that when *s* ≪ 1, scaling s by 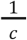 and *L* by *c* leaves *s*_*y*_ unchanged. Intuitively this makes sense: when the average number of offspring per mutant individual or *s* is reduced, but the number of generations *L* is increased or equivalently the timing between generations is decreased, then the percent growth per unit time remains unchanged. This is shown as a cartoon in Figure 2 where the viruses produce more children on average than the bacteria, but the timing between viral replication events is longer so that fitness of both populations is the same.

**Figure 2.**
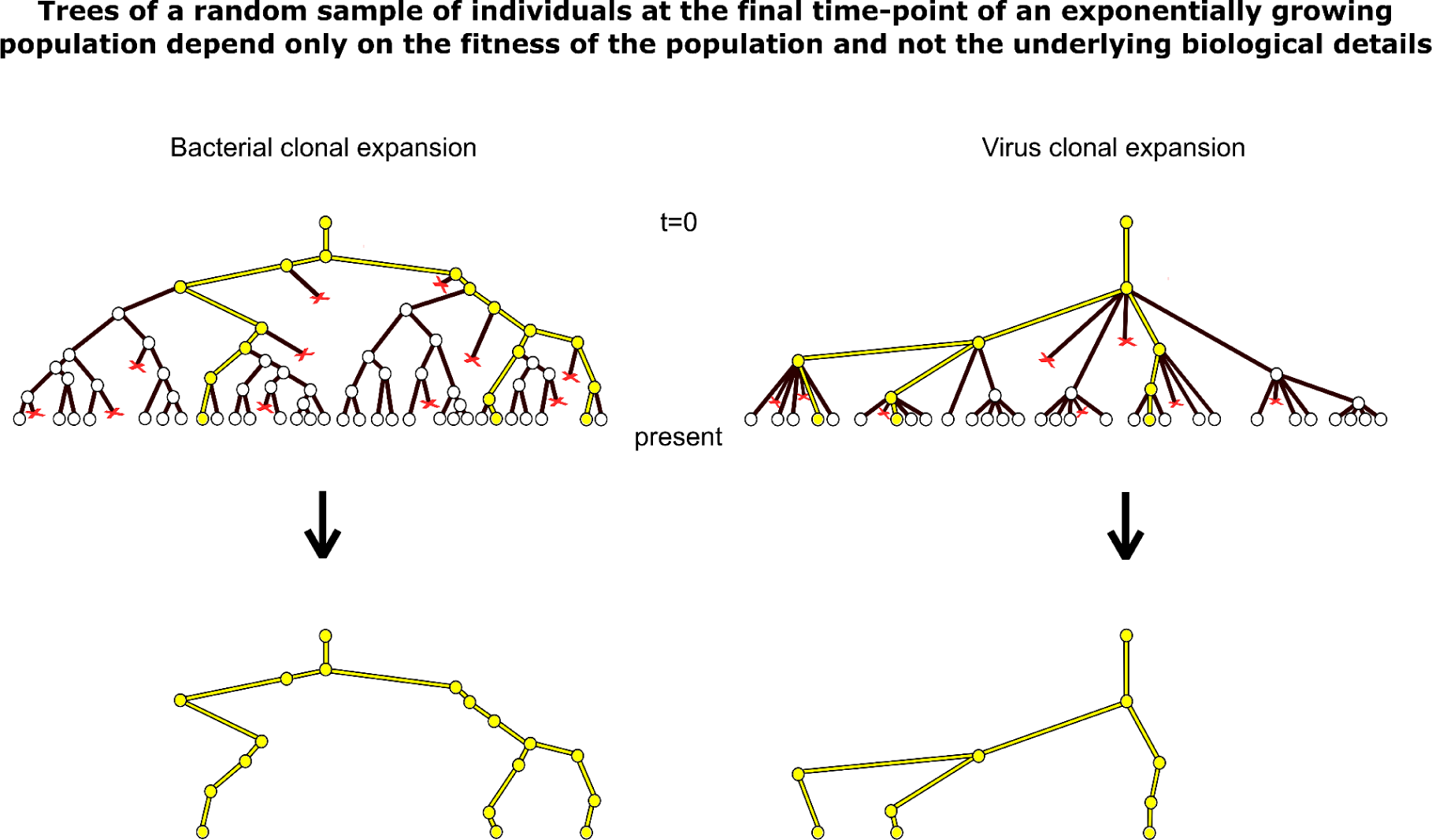
The schematic of a genealogical tree of a bacterial cell clonal expansion (left) and a viral clonal expansion (right) are shown side by side. The top node of the trees on the left and right represent the first cell and viral particle respectively that subsequently undergo replication events where children are produced, shown as nodes of the tree that branch off, and death events, shown as terminal red x marks. Time occurs from the top of the tree to the bottom of the tree in units of years, and only occurs in the vertical direction so that the time between any two points of the tree is their vertical distance. Reproduction events occur at a faster rate for the bacterial clonal expansion, but more children are produced at each replication event in the virus clonal expansion, and so the percent growth per year or fitness of both expansions is identical. A small number of cells is randomly sampled at the final time-point and their lineage tree is shown. Because both processes have the same fitness, the lineage trees look the same.

We then consider a clonal expansion with parameter values *s* and *L*, and then another clonal expansion with parameter values 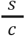 and *cL* so that *s*_*y*_ remains the same for both clonal expansions. In the first clonal expansion, the selective advantage is *s* and so the number of mutant individuals must fluctuate to 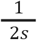 to escape drift and grow deterministically and exponentially [1], [9]. Under the rescaled parameters, the selective advantage is smaller, 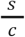, and since the trajectories are more neutral and drift more, the number of mutant individuals must fluctuate to a larger population size of 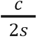 to escape drift and grow deterministically. As a result, the average number of mutant individuals through time is increased by a factor of *c* (Figure 3). Increasing the number of mutant individuals decreases the rate of coalescence since the probability that randomly selected lineages are closely related at each generation decreases. However, increasing the number of generations from L to cL also increases the rate of coalescence since there are more opportunities for lineages to coalesce per unit time [7], [11]. Taken together, these effects cancel and produce indistinguishable trees. Therefore, fitness determines the statistics of coalescence of lineage trees independent of the number of generations or equivalently the mutation rate per generation.

**Figure 3.**
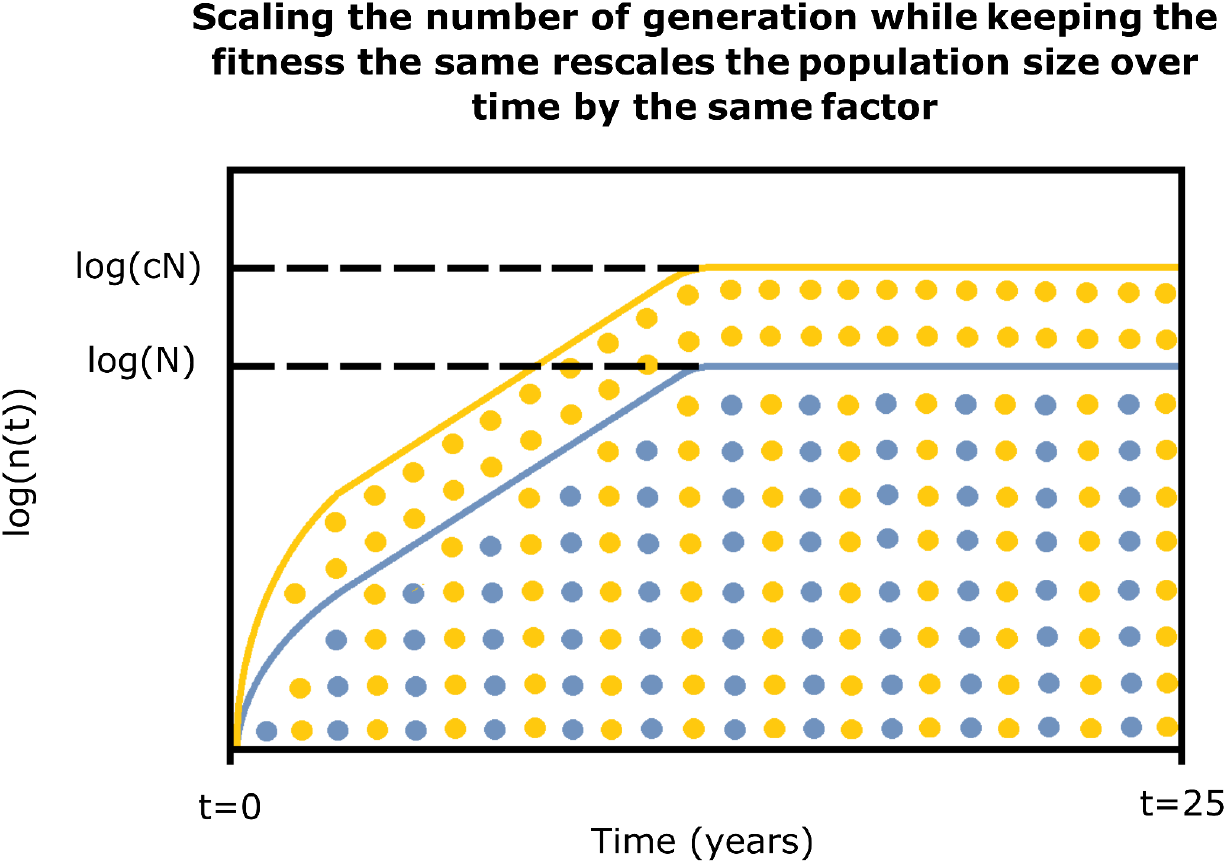
A schematic diagram showing how the number of individuals through time of a clonal expansion is scaled in the same way as the number of generations after escaping drift. The blue dots and line alone, ignoring the orange, is a schematic of the number of individuals through time of a characteristic clonal expansion. The blue line is just a trajectory on a log scale plotted against the time in years, and the blue dots represent the individuals at each generation and are not drawn to scale. Since the horizontal axis denotes time in years, and each vertical slice of blue dots represents a generation, then the spacing between the vertical slices of dots informs us about the timing between generations. If we now consider a clonal expansion where the timing between generations has decreased by a factor of two, denoted by the blue and orange dots combined, but the percent growth per year or fitness is kept the same, then the selective advantage decreases and the population size required to initially escape drift increases by a factor of two. This scales the population size through time by a factor of two, represented by the orange line. In summary, the population size through time is scaled by the same factor as the number of generations.

The intuitive arguments made in this section fall short of establishing the statistical invariance of trees for a given fitness. This is because our argument was based on the average size of the mutant clone. We did not account for the fact that there are fluctuations in the number of mutant individuals over time, and so every realization of a clonal expansion will have a different trajectory. These fluctuations in the number of mutant individuals mostly occur in the early history when drift dominates before the growth is deterministic and exponential. In particular, the clone of mutant individuals escapes drift when the number of mutants is ∼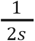 and takes on average ∼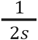 generations to do so, but the time it takes to escape drift is a fluctuating quantity [9]. Importantly, the trajectories do not converge to the average trajectory after a sufficient number of generations but always retain the variability inherited from the drift phase. This variability in the set of possible trajectories means that the coalescent statistics of trees must be computed across the set of all possible trajectories and cannot be computed using the average number of mutant individuals over time.

### Proof that fitness is invariant to the assumed number of generations

In the previous section we argued that when the number of generations is scaled by a factor of *c* in the Wright-Fisher model with the fitness (growth rate per unit time) kept constant, then the average number of mutant individuals over time will also be scaled by a factor of *c* so that the lineage trees will have the same coalescent statistics. This argument ignores the fluctuations in the number of mutant individuals over time that can also impact the coalescent statistics. To show that the invariance of coalescent statistics holds, we need to extend our argument from the average trajectory of the mutant clone size to all possible trajectories. Namely, we need to show that each possible trajectory that can be generated by the parameter values *s* and *L* will be scaled by a factor of *c* when using the parameter values 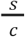 and *cL*, and that the scaled trajectory occurs with the same probability as the original trajectory. Although we do not have a way of specifying the distribution of trajectories in a simple form, we can specify the distribution of the number of mutant individuals at each time-point using Kimura’s diffusion approximation to the Wright-Fisher model with selection [10]. We will then use this distribution to prove the invariance of the coalescent statistics across the set of possible trajectories.

Kimura’s diffusion approximation is a version of the Fokker-Planck equation with a solution that gives the probability density φ(*x, t*), at any generation *t* of the frequency of mutant individuals *x* within a population of *N* individuals. If *s* is the selective advantage of mutant individuals and σ^2^ is the variance in the number of offsprings produced by each mutant individual per generation, then the diffusion approximation to the Wright-Fisher model with selection is [10]

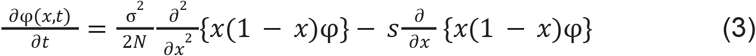

In the above PDE, *t* is expressed in number of generations, and σ^2^ = 1 for the Wright-Fisher model. However, to show our results, we first need to convert *t* to units of time. If the age of the population in units of time (such as years) is *a*, and *L* is the total number of generations, then the conversion from generations to time is given by the differential 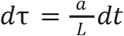. We then use the chain rule along with the differential to obtain:

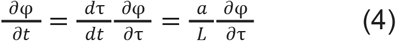

Plugging back into the PDE and multiplying both sides of the equation by 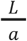 gives us

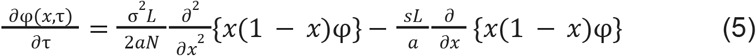

It is easy to see that when we replace the parameter values *s, L, N* with 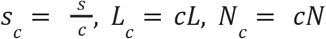 above, then the PDE and hence its solution or the frequency distribution over time remains unchanged. We scale the total population size N also by a factor c so that while the frequency of the mutant individuals is unchanged, the number of mutant individuals is scaled by c as expected. It is also clear from this scaling that the fitness, or percent growth per unit time, of the clone remains unchanged (refer back to the expression we derived for *s*_*y*_).

Importantly, scaling only *s, L* will not leave the PDE invariant. In this case, although the term that gives rise to the exponential growth (second term on the right-hand side of the equation) will remain unchanged, the drift term (first term on the right-hand side) will increase by a factor of *c* and will require a larger total population size *cN* to obtain cancellation. The way to interpret this scaling is that scaling the total number of generations *L* will increase the degree of fluctuations in the growing clone as a result of the decrease in *s* (the expansion dynamics become effectively more neutral per generation). Scaling *N* by *c* counteracts the increase in fluctuations because scaling *N* also increases the number of mutated individuals associated with any x at each time-point by a factor of *c*, which decreases the fluctuations due to drift. Taken together, the decrease in fluctuations due to the larger population size cancels the increase in fluctuations from scaling s.

A naïve argument might immediately suggest that fitness can be inferred without knowledge of the total number of generations *L* or equivalently the mutation rate per generation. Scaling the parameter values in the equation in the way we described leaves the PDE and hence the distribution of the frequency of the mutant clone over time unchanged. Since the size of the mutated population through time is scaled by a factor of *c*, and since *L* is also scaled by a factor of *c* then the coalescent statistics of the lineage tree should not change. However, this argument incorrectly assumes that the initial frequency does not change. In our model, we know that the clonal expansion begins with a single mutant individual, which corresponds to an initial frequency of 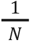 at the time at which the mutation occurs. Under this initial condition, if we scale the parameter values in the way described, the PDE remains invariant but the initial condition changes which changes the solution.

To correctly account for how changing the initial conditions changes the solution of the Fokker-Planck equation, we first need to condition the solutions on trajectories that do not go extinct. This is because when calculating the coalescent statistics, we are implicitly assuming that the mutant clone has survived extinction so that a lineage tree could be reconstructed at the final-time point. First, we need to make this assumption explicit in the Fokker-Planck formalism.

To do this, we define the random variable *X*_*t*_ as the frequency of mutant individuals as a function of time. The solution to the Kimura diffusion (Fokker-Planck) approximation would give the distribution of *X*_*t*_ at each *t*. We then relate the distribution of *X*_*t*_ conditioned on surviving extinction to the unconditional *X*_*t*_. As done earlier, define the unconditional frequency distribution of the mutant clone as φ(*x, t*). Then define *F* = {*fixation will occur*} to be the set of trajectories that have escaped stochastic extinction. The distribution of the conditioned *X*_*t*_ which is φ(*x, t* | *F*) can then be rewritten as

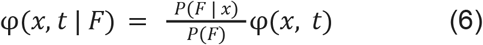

where we have used Bayes’ Theorem and have set *P*(*F* | *x, t*) = *P*(*F* | *x*), since the probability of fixation only depends on the clone frequency.

The probability of fixation of a clone with frequency *x* within a sufficiently large population of size *N* and with fitness 1 + *s* is derived from Kimura’s diffusion approximation [12]. We therefore have

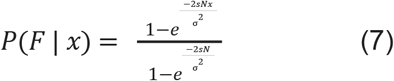

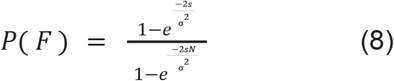

The probability of fixation *P*(*F*) independent of the clone frequency is simply the probability that a clone with frequency 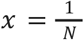 will eventually fix, since we always begin with one mutated individual. Substituting both probabilities back into the density function expression gives us

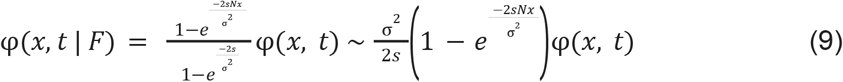

where we have assumed that 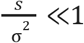. The unconditional gene frequency distribution φ is then given as the solution to Kimura’s diffusion approximation [10]:

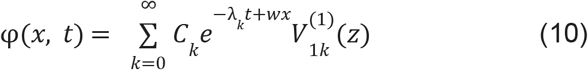

where

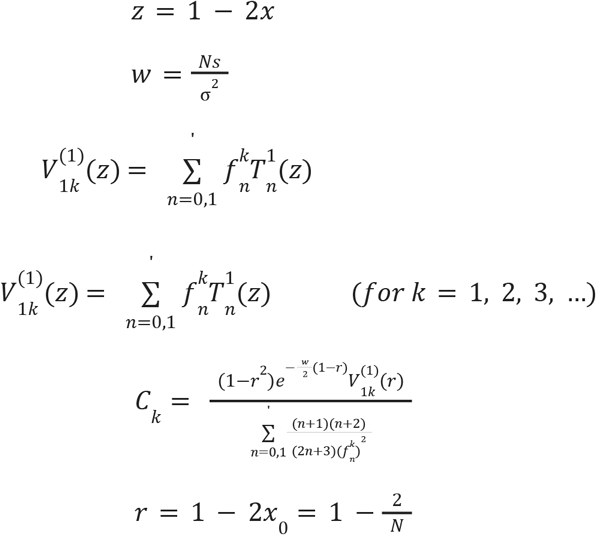

In the above equation, 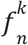 are constants, 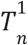 are Gegenbauer polynomials, and λ_*k*_ are eigenvalues [10]. The primed summation is over even values of n whenever k is even, and odd values of n whenever k is odd.

We then rewrite *C*_*k*_ in a way that will allow us to ignore second order terms:

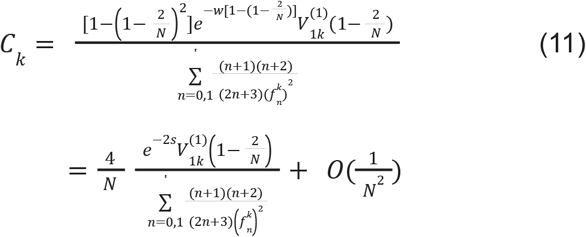

If we let *N*≫1 and 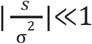 in the above expression, then since 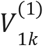 is differentiable and thus continuous (see time independent ODE for 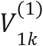 below) on the interval (−1, 1), and also has finite boundary conditions which are imposed to obtain the correct solution, we can continuously extend 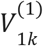 at *z* = 1 to obtain

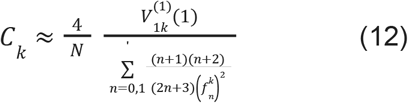

Then define 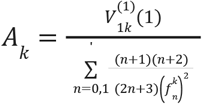, so that

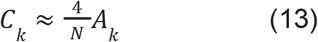

Substituting *C*_*k*_ into φ(*x, t*), and then φ(*x, t*) into φ(*x, t* | *F*) and 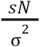 for *w* gives us:

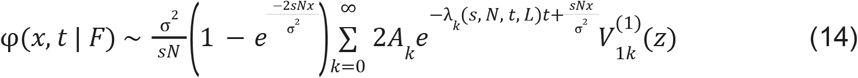

where we have changed the notation of λ_*k*_ to explicitly show the dependence of λ_*k*_ with respect to the parameter values we will scale. By substituting 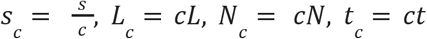, *L*_*c*_ = *cL, N*_*c*_ = *cN, t*_*c*_ = *ct* for *s, L, N, t* in φ, it is easy to see that φ remains unchanged whenever 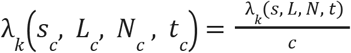.

To show that 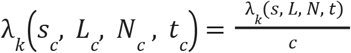, we first note that the unconditional solution to φ is a linear combination of solutions 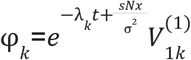 (refer back to Eq. 10). When plugged back into the Fokker-Planck equation, we get an ODE for each 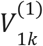 and its corresponding λ_*k*_ :

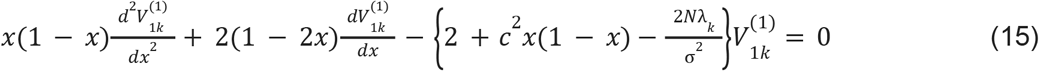

For a given set of parameter values *s, L, N, t*, there are infinitely many solutions to the above ODE, one for each eigenvalue λ_*k*_. If we rescale the parameters as before, then 2*N*λ_*k*_ increases by a constant factor of *c*. After rescaling, the eigenvalues λ_*k*_ in the equation must be divided by a factor of *c* to be associated with the same solution 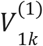. Therefore, for the parameter values 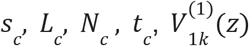 is also a solution with eigenvalue 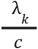. The invariance of φ(*x, t* | *F*) then follows from the fact that 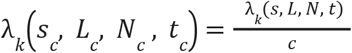, and from the fact that the scaled eigenvalue is associated with the same 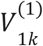 so that each 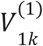 is unchanged in the solution.

Therefore, we have shown that the distribution of *X*_*t*_ conditioned on survival remains unchanged when the parameters are scaled, and thus the frequency of the mutant clone over time remains unchanged. However, since *N* is one of the parameters being scaled and the frequencies over time remain the same, then the number of mutant individuals at every time-point is scaled by a factor of *c*. Scaling the size of the population of the mutant clone by the same factor as the number of generations leaves the coalescent statistics unchanged.

Taken together, we have shown that the coalescent statistics of lineage trees do not change when the number of generations *L* and population size *N* are scaled by *c*, and the growth rate per generation *s* is scaled by 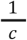. Importantly, this scaling does not change the growth rate per unit time or fitness *s*_*y*_. Conversely, this conclusion implies that we can infer the fitness from observed lineage trees by assuming an arbitrary mutation rate to convert the branch lengths in number of mutations to number of generations. The inferred number of mutant individuals will also be scaled by an arbitrary number that depends on the assumed mutation rate; however, the inferred fitness will match its true value.

To complete the proof, we note that in the above derivation we only showed that the distribution of the frequency of the mutant clone at every time-point is invariant to the proposed scaling. This does not necessarily imply that the frequency of the mutant clone as a function of time (referred to as the trajectory) is also invariant. To show this, we write down a stochastic differential equation for the frequency of the mutant clone [13]:

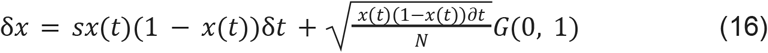

Here *x* is the frequency in number of mutant individuals, *t* is time in number of generations, *x*(*t*) is the value of *x* at time *t, N* is the total number of individuals, and *G*(0, 1) is a Gaussian distribution with mean zero and variance one. We can then convert time from number of generations to years using the differential 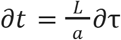 and obtain

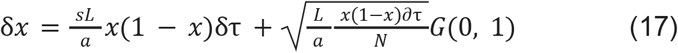

If we scale the parameter values of this equation as before by replacing *s, L, N* with 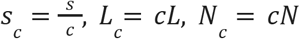, the equation remains unchanged provided we begin with the same initial frequency. This remains true even if we condition the equation on no stochastic extinction. However, as before if we begin with an initial number of one mutant individual then the initial condition of 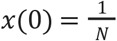 changes when we scale the parameters. To circumvent this problem we can use the Fokker-Planck equation conditioned on surviving extinction. Previously in this section, we showed that the distribution of frequencies at each time-point conditioned on survival remains unchanged under the proposed scaling, even when we begin with an initial condition of one mutant individual. This invariance breaks down near generation *t* = 0 when the initial frequencies differ by a factor of *c*, but holds true for all generations *t* > ε. Although we do not have a way of computing ε directly, as described in the next section, we can bound the error on the inferred time at which the first individual of a population arose by relating ε to its effect on the coalescent structure in the early history of the expansion.

Taken together, above results show that the coalescent statistics of phylogenetic trees are invariant when model parameters are scaled from *s, L, N* to 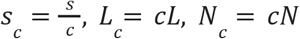. Importantly, the fitness remains constant under this scaling (for an intuitive explanation for why *N* must also be scaled even though fitness is independent of *N*, see Figure 4). Therefore, if fitness is inferred by fitting the model to an observed phylogenetic tree, the correct value of fitness can be obtained even if the duration of each generation was chosen arbitrarily and only correct up to a scaling with respect to the true value of the underlying biological process that generated the observed tree.

**Figure 4.**
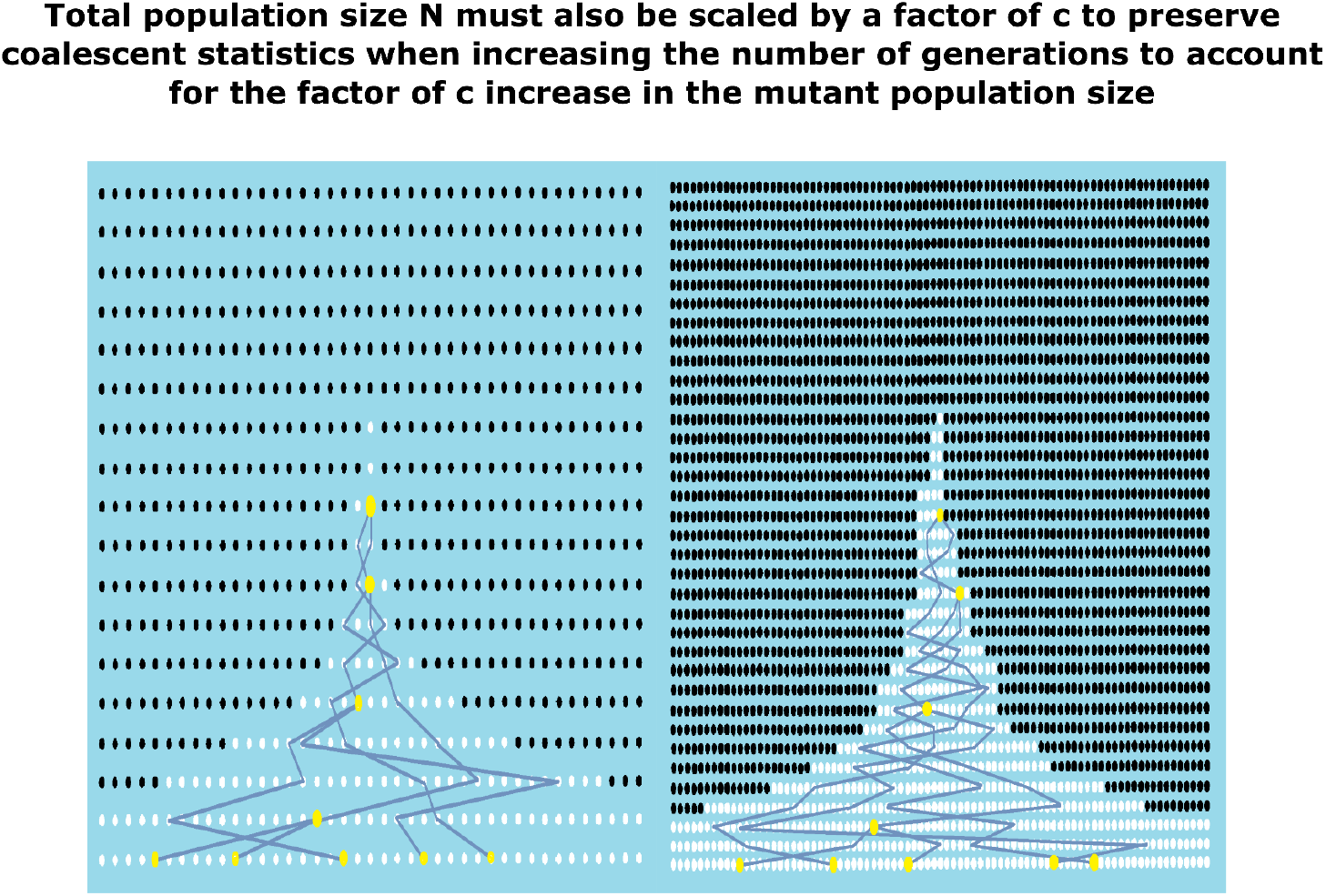
Schematic shows why *N* must also be scaled by a factor of *c* when scaling *s* and *L* by a factor of *c*. When a clonal expansion is generated under the scaled parameter values 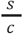 and *cL*, the number of mutant individuals over time increases by a factor of *c* and so *N* must be scaled by a factor of *c* so that saturation is reached at the same time as the unscaled process which is necessary to preserve the coalescence statistics. The left hand side shows the unscaled process and the right hand side shows the process with scaled parameter values. The white dots are the mutant individuals, and the yellow dots show the point of coalescence of the trees which are similar in both figures.

### The error in the inferred time at which the first mutant individual arose is at most one generation

Above we showed that fitness can be inferred without knowledge of the mutation rate per generation or equivalently without knowledge of the duration of each generation. In addition to fitness, for many applications, we would like to infer the time at which the first mutated individual arose. For example, we would like to infer the age at which the cancer mutation first occurred in a patient from the phylogenetic tree of individual cancer cells obtained from the patient. Here, we will show that the time at which the first mutant individual arose (time of onset) can be inferred from the phylogenetic tree without knowledge of the duration of each generation. In particular, we show that the inferred time of onset is different from the true time of onset on average by at most the duration of one generation. We will show this first for the simpler case of the Wright-Fisher model with σ^2^ = 1, where σ^2^ is the variance in the number of offspring per individual, and then generalize the results to any value of σ^2^. More formally, we will show that if the time between subsequent generations in the model is assumed to be *T*_*c*_ and the true time between subsequent generations is *T*, then the mean of the distribution of inferred time of onset is shifted into the past by at most *T* − *T*_*c*_ if *T* > *T*_*c*_ and into the present by at most *T*_*c*_ − *T* if *T* < *T*_*c*_. For example, if we assume that an expanding population of cancer cells divides once per year in the model, then the inferred time of onset will be shifted to the past by at most one year on average if the cancer cells divide at a rate of once every two years and by half a year into the present if they divide twice per year. To understand where these bounds come from, we first note that when a population is expanding according to the Wright-Fisher model and is in the drift phase, the average population size grows linearly by one individual per generation when conditioned on surviving extinction [13]. We then consider trajectories of the mutant population size *x*(*t*) (expressed in terms of frequency, or fraction of the total population) and also trajectories *x*_*c*_(*t*) where we have rescaled the parameters describing the population growth by a factor of *c* ≥ 1, replacing *s, L, N* with 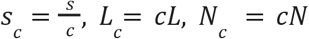 as the duration of each generation for *x*(*t*) and *x*_*c*_(*t*) respectively. For each generation of the original process, *c* generations pass in the scaled process, and hence 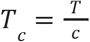. As described in the previous section, the original trajectory and the scaled trajectory eventually become statistically identical (See Figure 5 to see how both trajectories differ by approximately a factor of *c* even during the linear drift phase. This is equivalent to their frequencies being identical since their total population sizes differ by a factor of *c*). However, because both trajectories must start with one individual at time *t* = 0 when the first mutant individual arose, or since 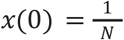 and 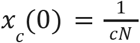, they cannot be identical for *t* < ε sufficiently close to *t* = 0. Since both trajectories grow linearly with the same slope per unit time during drift, and since *x*_*c*_(*t*) begins at a smaller frequency, we can get the average values of the original and scaled trajectories to match exactly if we start the scaled trajectory earlier by some Δ*T*. We can then find Δ*T* such that 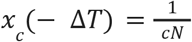, and such that by time 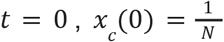. Δ*T* satisfies 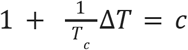, which we can use to obtain 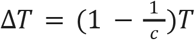. Using the definition of *T*_*c*_, it follows that the required offset for the average of two trajectories to agree is *T* − *T*_*c*_. This argument only holds for *c* ≥ 1. For *c* < 1 the required shift is *T*_*c*_ − *T* into the present using a similar argument that shifts the start of the original trajectory backwards in time with respect to the scaled trajectory.

**Figure 5.**
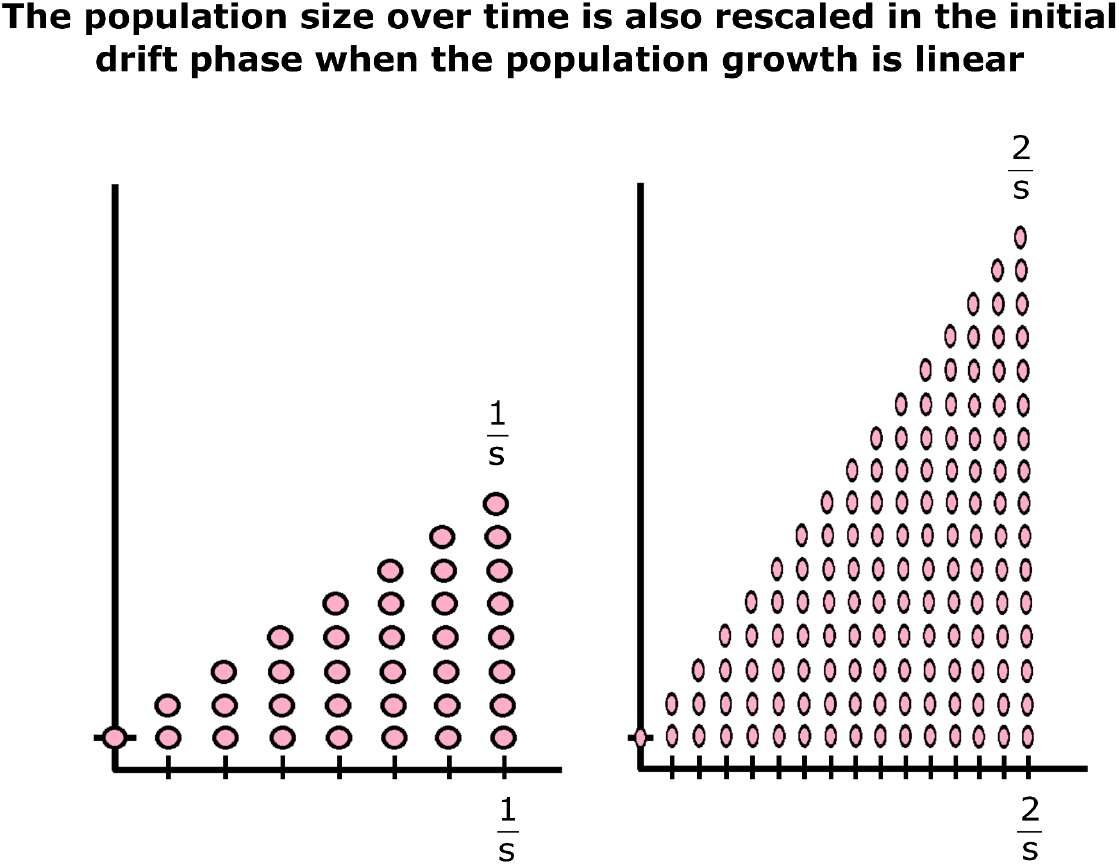
Schematic plots show how the number of individuals as a function of time, shown in red circles, is scaled in the drift phase for two clonal expansions that differ in the timing between generations but where the percent growth per year or fitness is the same after the growth of the clones become deterministic and exponential. The vertical axis is n(t), the number of individuals, and the horizontal axis is time in years, but tick marks to denote the number of generations per unit time are superimposed to show how the timing between generations differ across both plots. On the right hand side the timing between generations is shorter by a factor of two. Since the percent growth per year or fitness is the same for both processes after escaping extinction, and since the selective advantage is smaller by a factor of two for the process plotted on the right because the number of generations is larger, then the number of generations required to escape drift is twice as large keeping the time in years to escape drift the same across both plots. However, the decrease in selective advantage also increases the population required to escape drift by a factor of two.

Although the offset introduced above results in perfect agreement between the means of the original and scaled trajectories, it does not do so for all the higher moments. In fact, all higher moments *x*(*t*) are exactly equal to that of *x*_*c*_(*t*) without introducing any offset. To show this, we will show that conditioned on surviving extinction, the higher order *k*^*th*^ centered moments during drift in terms of frequency are

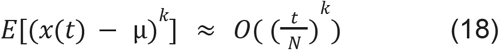

We can then replace *N, t* with *N*_*c*_ = *cN, t*_*c*_ = *ct* in the expression above to show that the higher moments do not change even when *s* is replaced with 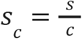 such that the fitness remains the same. As a result, the fluctuations in the frequency of mutant individuals agree perfectly when no offset is introduced. Therefore, we can either introduce an offset between the original trajectory and the scaled trajectory to bring their means into agreement or introduce no offset and have all their higher moments in agreement. Any inference framework will use a combination of the moments of the trajectories, obtained indirectly from the coalescent statistics of the reconstructed phylogenetic tree, to infer the time of onset. Therefore, the inferred time of onset will be different from the true time of onset by at most the required offset to align the means at the expense of aligning the higher moments.

To complete our argument, we now derive the expression for the moments conditioned on surviving extinction. We first derive the moments without conditioning on surviving extinction and then relate the conditional moments to higher order unconditional moments.

Since the dynamics of clonal expansions break down only in the early history, we can ignore the deterministic term in the stochastic differential equation, include σ^2^ so that our results are more general, and rewrite it in a modified form to model the drift phase as:

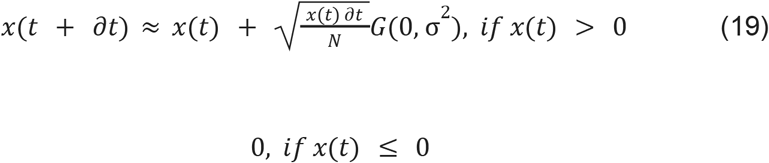

Since *x*(*t* + ∂*t*) is a function of the random variables *x*(*t*) and *G*(0, σ^2^), and since *x*(*t*) and *G*(0, σ^2^) are independent and 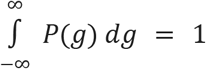 where *P*(*g*) is the density of *G*(0, σ^2^), we can compute the expectation value as:

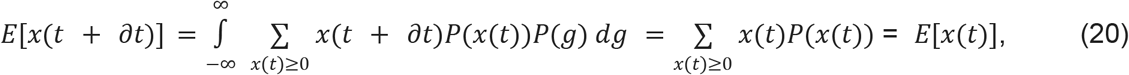

where the second equality is justified by replacing *x*(*t* + ∂*t*) for the cases *x*(*t*) > 0 and *x*(*t*) = 0.

By recursing and then using the initial condition 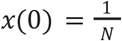 we obtain:

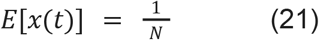

To obtain the unconditioned second moment, we first square both sides of the stochastic differential equation:

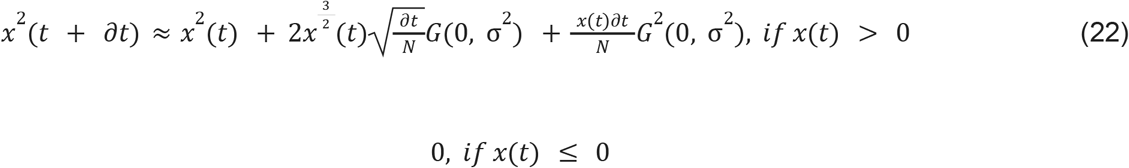

We then compute the expectation value in a similar manner as before by remembering to replace *x*^2^(*t* + ∂*t*) for both cases *x*(*t*) > 0 and *x*(*t*) = 0, except in addition we use 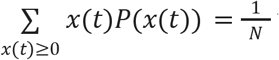 which is the expectation we previously computed, 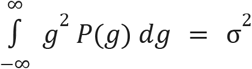 because the expectation of *G*^2^ (0, σ^2^) is the second moment of a Gaussian with mean zero, and 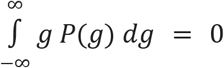 which causes the middle term to vanish:

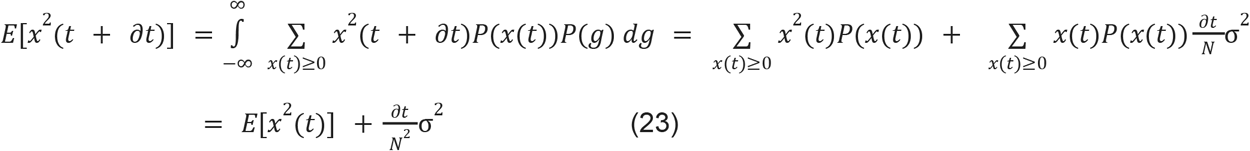

We then solve the recursion with an initial condition of 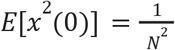 to obtain:

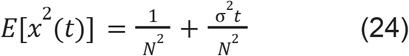

A similar procedure can be used to compute the third moment and the higher order moments too:

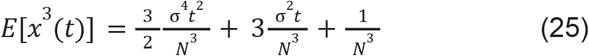

We can then relate any *k*^*th*^ conditional moment to the unconditional (*k* + 1)^*th*^ moment. This is done by first defining *S* to be the conditioning event of the set of all trajectories that escape extinction which depends on the parameter *s*, and then computing

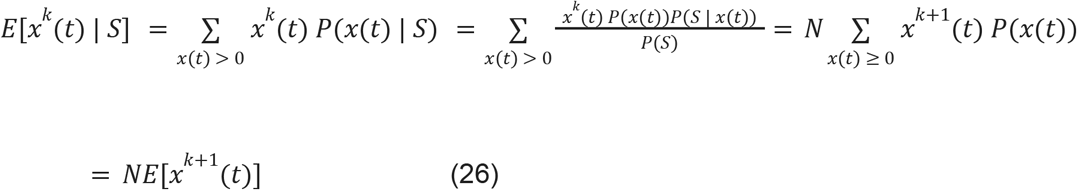

where we have used Bayes’s Theorem and in the third equality the probabilities of surviving extinction 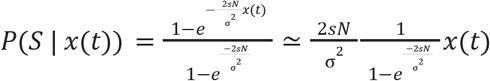 and 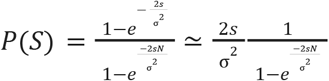 assuming *s ≪* 1 [12], and changed the sum to include *x*(*t*) = 0.

Using this expression and the unconditional moments we can then derive:

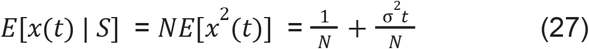

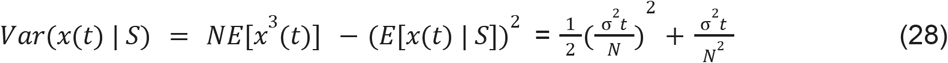

By iterating this procedure, we can then show that all the centered moments are 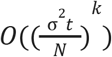.

Simulations that validate this result are shown in Figure 6a-c and the details of computation are discussed in the later section titled ‘Centered moments of simulated expansions match their theoretical values.’ Figure 7 shows the impact of an incorrect number of generations on the inferred time of onset from simulated data and the details are found in the later section titled ‘The impact on the inferred age of onset when ABC model has incorrect assumptions’.

**Figure 6.**
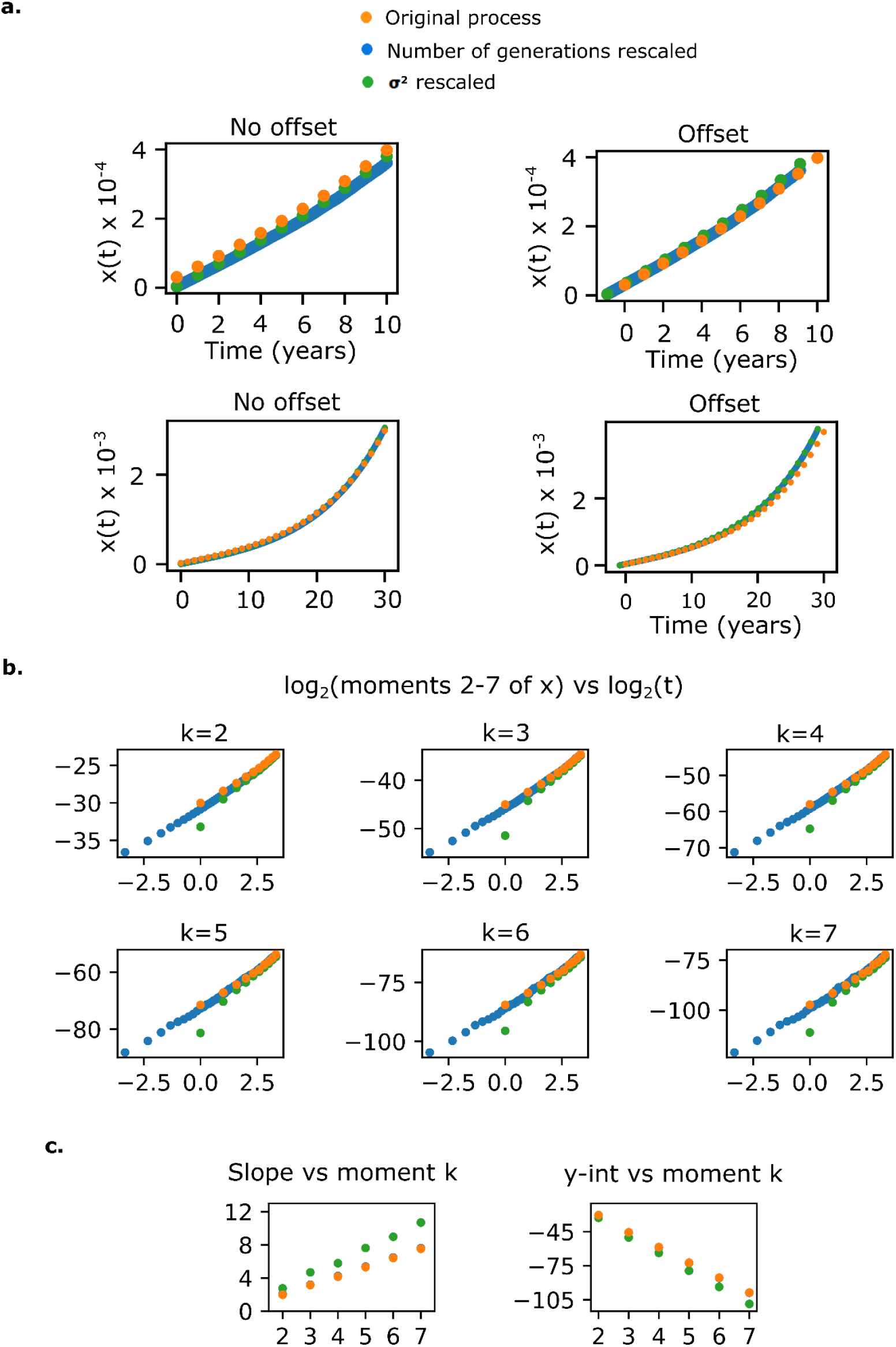
To show the k-th centered moments grow as 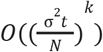 in the initial drift phase, 10,000 clonal expansions with fitness *s*_*y*_ = 0. 1 were simulated according to the Wright-Fisher process with selection for 200 generations and conditioned on surviving extinction at the final time-point (orange dots). Another set of 10,000 clonal expansions were simulated using 10 times as many generations per year but keeping the same fitness value (blue dots). A third set was simulated under the same parameters as the first set except that only 10% of the population was retained at each generation (green dots). The total population size *N* was scaled by a factor of 10 in the last two simulations to keep the extent of drift the same across all three simulations. **a)** Offsetting the scaled process aligns the means initially. Plot on the top left hand side shows the mean frequency x as a fraction of *N* over time in years across the 10, 000 simulated expansions. Since all three processes begin with 1 individual, and the latter two processes have a larger *N*, their frequencies differ and the means do not line up initially. However, if we offset the processes by the predicted value 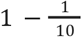, the means perfectly coincide and are shown on the top right. The bottom two plots are identical to the top two except the mean values are shown for a longer period of time. As predicted, the means will coincide eventually even if their frequencies initially differ shown on the bottom left. On the bottom right, we see that even though initially we can get the means to match, the mean trajectories will eventually diverge. **b)** *k* ^*th*^ centered moments grow as O(*t*^*k*^) initially. *log*_2_ (*k centered moment of x*) vs *log*_2_ (*t*) plots for the centered moments 2-7 of x for the same expansions. Consistent with the predictions that the plots should be linear and that the plotted values should coincide, the centered moments match well with no offset introduced except for a slight deviation from the expected behavior in the initial generations for the expansions that underwent large sequential bottlenecks (green dots). **c)** For each of the plots in b), a line was fit for each of the three processes (blue dots covered by orange dots because they match perfectly) and the slopes and y-intercepts plotted as a function of k on the left and right respectively. The slight deviations of the green dots is likely due to the initial deviation at *log*_2_ (*t*) = 0 in the plots in b).

**Figure 7.**
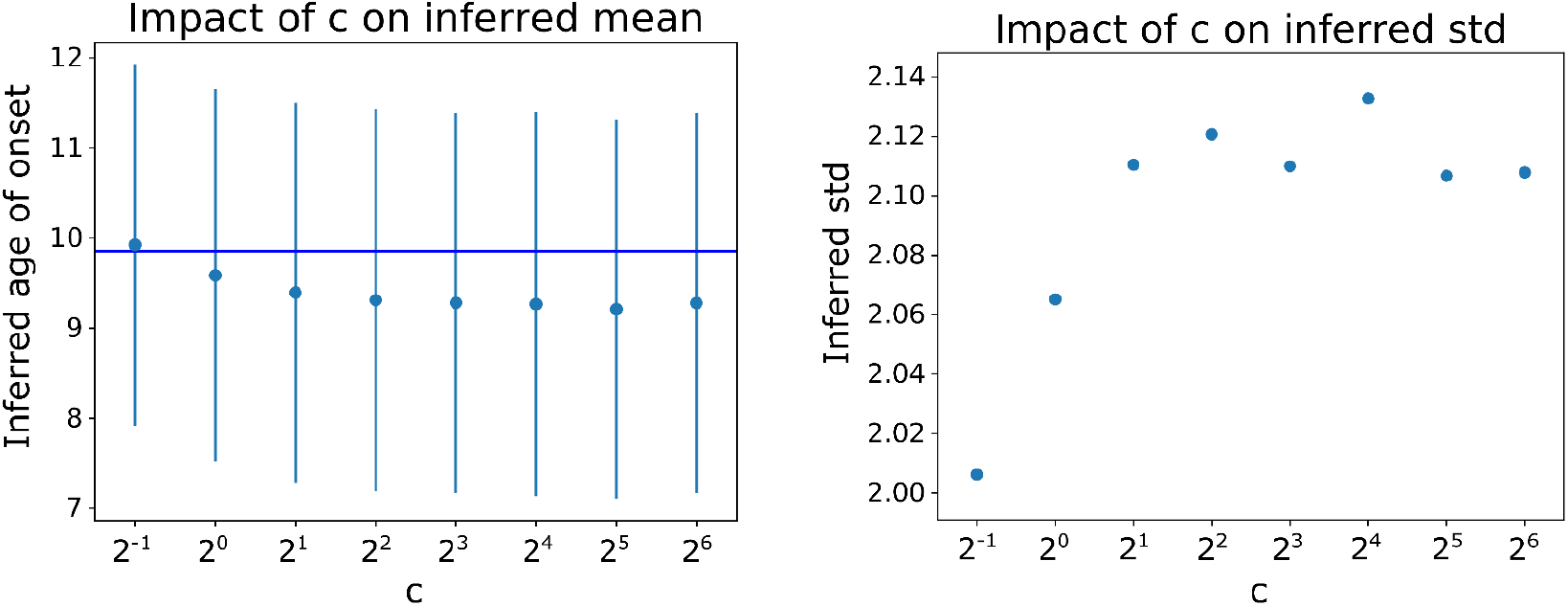
Inferred age of onset is shifted by at most the duration of one generation. ABC inferences were carried out repeatedly on the same simulated data tree constructed from a simulated cancer for a 34 year old patient while varying the number of generations assumed in the ABC model. The mean of the inferred age of onset is plotted for each ABC inference against the particular value of *c*, where the model used for ABC inference assumed *c* times as many generations as the true number of generations in the data tree (which was 2 generations per year). Error bars denote 1 std, and the exact value of each std is plotted against *c* on the right hand side. As expected, as the value of *c* increases, the error in the inferred age of onset approaches the theoretical bound of 1 generation.

Finally, the arguments made in this section can be easily extended to arbitrary values of σ^2^. This is because more generally the population size increases by σ^2^ at each generation on average during drift as opposed to one (when we assume σ^2^ = 1). Therefore, more generally, the inferred time of onset is off by at most the time that it takes the mutant population to increase in size by one individual. This time can be shorter than one generation if σ^2^ > 1 in the underlying process. In the next section, we will consider the implications on the inferred time of onset and fitness if the true value of σ^2^ itself is not known.

### Coalescent statistics do not depend on the specific distribution of the number of offspring or fluctuations in the timing between generations

The Wright-Fisher model assumes that the distribution of the number of offspring generated by each individual in the population at each generation takes on a specific form. It also assumes discrete and fixed timing between generations across all lineages. Our results would not be very interesting if our derivations only held for the Wright-Fisher process, which is a very specific model that is unlikely to be correct in most applications. For example, the specific distribution of the number of offspring may be unknown or could exhibit large fluctuations or functional forms not well approximated by the Wright-Fisher process, and there may be additional fluctuations in the timing between replication events across the lineages. We therefore set out to show in the following section that the coalescent statistics of lineage trees are independent of the distribution of the number of offspring per individual which is also equivalent to incorporating fluctuations in the timing between generations (see Figure 8)

**Figure 8.**
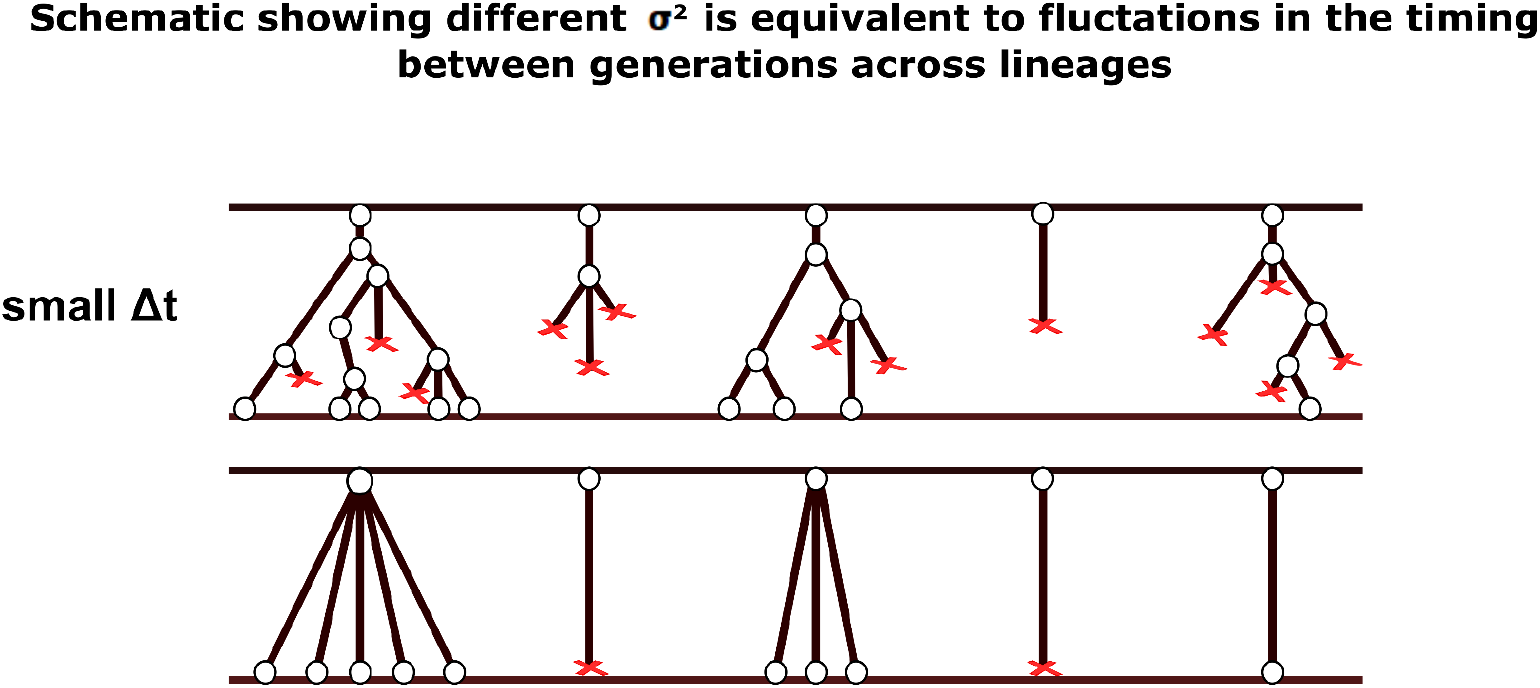
Schematic showing how changing the variance of the number of offsprings produced by each individual every generation, σ^2^, is equivalent to introducing fluctuations in the time between generations across the lineages. The top figure shows a birth-death process where the number of generations fluctuates, and the bottom figure shows a different birth-death process with discrete generations. In the process illustrated at the top, we assume Δ*t* is the average time between 1 generation. The process shown on the bottom has discrete generations of duration Δ*t*. Since the distribution of the number of offspring produced is the same after time Δ*t* in both processes, these processes will produce similar coalescent statistics across many generations.

We can make an intuitive argument for why the coalescent statistics of lineage trees should not depend on the specific distribution of the number of offspring produced by each individual. When we scale the variance in the distribution of the number of offspring, we increase the amount of drift and so the minimum population size required to escape extinction is scaled by a similar amount. This implies that the average number of mutant individuals at each time-point is scaled by the same factor which decreases the rate of coalescence. However, if there is a higher variance in the number of offspring, then a smaller fraction of the mutant individuals are giving rise to the subsequent generation at any time point, which speeds up the rate of coalescence. These two opposing effects cancel leaving the coalescent statistics of the lineage trees unchanged.

To show this mathematically, we consider the Canning model [14], [15] which is similar to the Wright-Fisher model except that the distribution in the number of offspring produced by each mutant individual at each generation is arbitrary but identical across the mutant individuals, and has a variance σ^2^. We then present our derivation that the rate of coalescence scales proportionally to σ^2^ and inversely to *N*, and does not depend on the specific form of the distribution. We then revisit the equations in the previous sections to show that the frequency over time remains unchanged after scaling both parameters σ^2^ and *N* by the same factor of *c* which implies the population size as a function of time increases by a factor of *c*. Taken together, this shows that the increase in the rate of coalescence from scaling σ^2^ is canceled by the decrease in the rate of coalescence from the increasing population size. Since the fitness of the population does not change when scaling σ^2^ and *N* (recall fitness only depends on *s, L*, and *a*), and since the statistics of trees also do not change under this scaling for a given fitness value, then the fitness of a population can be inferred from lineage trees even if the assumed distribution in the number of offspring does not match the true distribution in the underlying process.

We now derive the coalescent rate per generation under an arbitrary distribution of the number of offspring per individual per generation with variance σ^2^, and with mean 1 + *s* so that the frequency of mutant individual grows by a factor of 1 + *s* per generation. We also assume that the number of mutant individuals is large so that the frequency grows deterministically, and that *s ≪* 1. We assume that there are *n* mutant individuals at generation *t*, and compute the rate of coalescence when the clone size grows from *n* at time *t* to *n*(1 + *s*) at time *t* + 1.

We define the probabilities of each mutant individual at generation *t* having *c* children as *p*_*c*_. Since *p*_*c*_ is just the fraction of individuals that have c children at generation *t*, then the probability we randomly sample at *t* + 1 an individual derived from a parent with *c* children is *cf*_*c*_, where 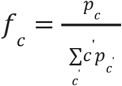. To compute the probability of coalescence at any generation *t* given *k* individuals at *t* + 1, we compute 1 minus the probability that no coalescence occurs. To determine the probability that no coalescence occurs across the k individuals, we begin by calculating the probability of the second individual not coalescing with the first individual. This is given by 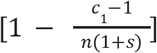 which is just the probability that the second individual does not share a parent with the first individual given that the first individual is the child of a parent with *c*_1_ children, and then averaging across all possible *c*_1_ (see Figure 9 to better understand the indices *c*_*i*_) to obtain

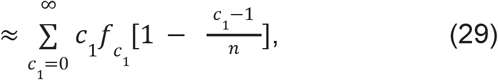

where we have assumed *s ≪* 1. To simplify the math, we will always replace 1 + *s* with 1 whenever it shows up at any step in the derivation because 1 + *s* will become 1 in the final answer anyways when we let *s ≪* 1.

**Figure 9.**
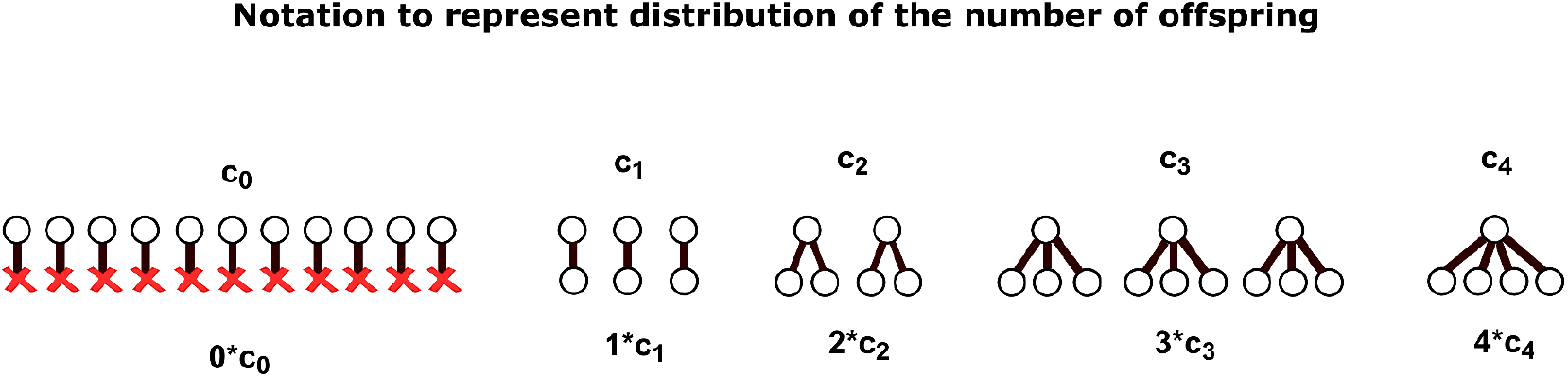
Schematic showing what each *c*_*i*_ represents in Eq. 29-31. In particular, *c*_*i*_ is the number of individuals in a given generation that produce *i* offspring in the next generation.

We then include the probability of the third individual not coalescing with individual 2 or individual 1, again averaging over all possible *c*_2_. This process is iterated up to individual k, resulting in the following nested sum

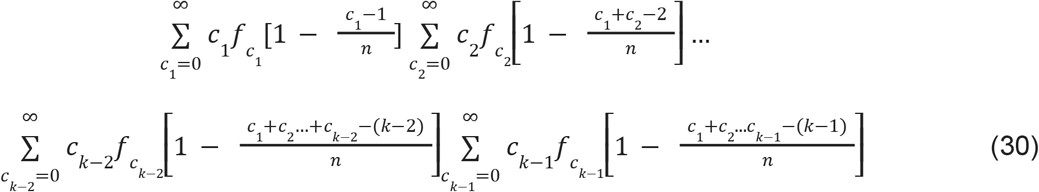

Note that we can distribute the innermost sum and use the equalities 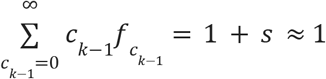 and 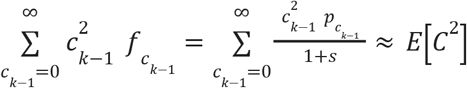 where *C* is the random variable denoting the number of offspring per individual per generation to obtain

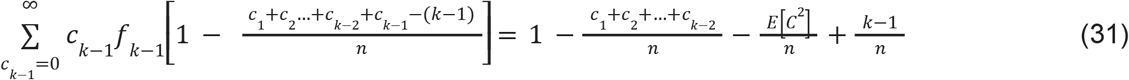

If we continue to iteratively multiply innermost terms followed by distributing the innermost sum and therefore working our way from the inside out, we can show that the sequence of nested sums is equal to

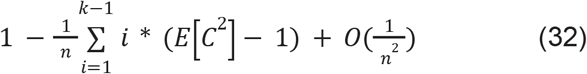

Then note that *E*[*C*^2^] = *Var*(*C*) + *E*^2^[*C*] = σ^2^ + (1 + *s*)^2^ ≈ σ^2^ + 1.

Substituting and letting second order terms vanish since *n* ≫ 1 gives us

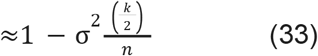

It then follows that the rate of coalescence per generation is

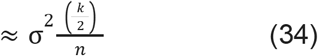

Comparing this to the classical Kingman coalescent probability, we observe that the variance σ^2^ in the number of offspring has effectively rescaled the population size of the mutant clone from n to 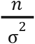 such that the coalescent rate has increased to the coalescent rate of a population under the Wright-Fisher model but with population size 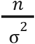. Increasing σ^2^ by a factor of *c* therefore effectively decreases *n* by a factor of *c* which requires a factor of *c* increase in *n* to preserve the rate of coalescence. We now revisit the equations previously derived to show that the mutant population size *n* is also scaled by a factor of *c*.

To show that the mutant population size scales by the same factor of *c* as σ^2^, we can follow the steps of the invariance proof identically except that we instead replace *N* and σ^2^ with *N*_*c*_ = *cN* and (σ^2^)_*c*_ = *c*σ^2^ in the Fokker-Planck equation to show that the frequency over time *x* is unchanged. Since *N* is scaled by a factor of *c, n* associated with any *x* is scaled by a factor of *c* as well implying that the mutant population size over time is also scaled by a factor of *c*. We can then rescale these same parameter values in the stochastic differential equation to show the invariance of individual trajectories as well.

Similar to scaling the number of generations, rescaling *N* and σ^2^ in the Fokker-Planck and the stochastic differential equation only show that the trajectories over time are scaled by a factor of *c* for *t* > ε, but are not scaled by a factor of *c* near *t* = 0 because there is an initial condition of one individual. If we account for the added variance in the number of offspring, we can show that the inferred time of offset is incorrect by at most the time it takes for the population size of the mutant clone to increase by 1. To show this we first note that the clone size grows as 1+ σ^2^ *t* on average, and the second and higher moments expressed as frequency of the total population size *N* are given by ∼ 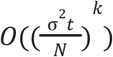 after conditioning on surviving extinction. Since σ^2^ and *t* are expressed as the product σ^2^ *t* in the moment expressions, it follows that scaling σ^2^ by *c* is equivalent to scaling the number of generations *t* by *c*, which is equivalent to scaling the *c* duration of a generation by 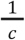. Intuitively, this makes sense because shortening the duration between the generations means that the clone grows more per unit time, which also happens when σ^2^ is increased. Therefore, as we argued in the previous section, we can get the average values of the original trajectories and the scaled trajectories that use the parameter (σ^2^)_*c*_ = *c*σ^2^ to match exactly if we begin the scaled trajectory earlier by the same amount of time required to align the average values of the original trajectory with a trajectory whose generation time has been scaled by by 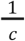. In the previous section, we showed that the inferred time at which the mutant population arose is different from its true value by at most one generation. Using a similar mathematical procedure and the equivalence of scaling σ^2^ with scaling time, we can obtain a more general expression that bounds the difference between the true time of onset and the inferred time of onset by the average time it takes the mutant population to increase by one individual.

### Coalescent statistics do not change if the fitness of each individual fluctuates

In the previous sections we had assumed that the fitness of each individual is identical. However, in most practical applications, the notion of fitness is a coarse-grained average growth rate which may vary from individual to individual and across generations. To show that heterogeneity in fitness does not change the coalescent statistics of lineage trees, we will assume that the fitness of each individual at each generation is chosen identically and independently from a fixed fitness distribution. We emphasize that the fitness of a parent is uncorrelated with that of its children. If differences in fitness are passed onto the next generation, the coalescent structure of lineage trees clearly changes as individuals with the highest fitness start to dominate the population (see [16], [3], [17]). To incorporate heterogeneity in fitness, we modify the Wright-Fisher model with selection as follows. As before, at each generation, individuals give birth to a new generation and immediately die out. Each individual in the new generation descends from a wild type individual with probability *p* and from the *i* ^*th*^ mutant with probability (1 + *s*_*i*_)*p* (for *i* = 1, …, *n*). Since probabilities must sum to 1, we have the condition:

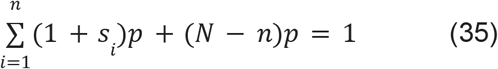

This means that the probability an individual at generation *t* + 1 descends from the *j*^*th*^ mutant individual is:

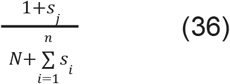

The number of times the *j*^*th*^ mutant is picked is therefore a binomial distribution with parameters N and 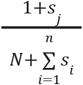, conditional on the values *s* which are i.i.d. random variables. If we assume that 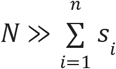, the expected number of children produce by the *i*^*th*^ mutant is 1 + *s*_*j*_, which means the dependence on all *s*_*i*_ ≠ *s*_*j*_ has vanished. Since 1 + *s*_*j*_ is the conditional mean number of children, then the unconditional mean number of children must be 1 + *s*_µ_, where *s*_µ_ is the mean of the distribution *S*.

Now recall that our results in the previous section show that the statistics of trees from a random sample at the final time-point do not depend on the distribution in the number of offspring per individual per generation and only on fitness. It then follows that the mean value of fitness can be inferred from the coalescent statistics of the lineage trees even if the fitness distribution is not known.

### Age of onset and fitness values of multiple clones competing within a population can be inferred simultaneously

Throughout this paper, we have made the simplifying assumption that lineage trees are constructed from randomly sampled individuals of an exponentially growing population where only a single, selectively advantageous mutation produced the exponential growth. This assumption ignores many realistic scenarios, such as when cancer cells acquire subsequent driver mutations that produce new subclones with higher fitness values, or when HIV infects cells for multiple generations and various mutations with differing fitness values arise in different lineages and compete with each other (i.e. clonal interference). We can extend our results to these more complex scenarios to show that the lineage trees constructed from randomly sampled individuals from populations that contain multiple competing mutations can be used to infer the history of expansion of multiple subclones and the times at which the subclones emerged without knowing the specific biological details of each expanding clone.

To illustrate this point, we can define a simple model of multiple driver mutations that builds off of the Wright-Fisher process with selection. The new model is identical to the Wright-Fisher process with selection we defined previously, except a secondary mutation with a different fitness advantage to the first arises in the population either in the same lineage, such as a cancer with a subsequent driver mutation where a subclone of the cancer cells is expanding more rapidly than the other cancer cells, or in a different lineage such as two strains of a virus that each have a distinct mutation and compete with each other. The only difference in the new model occurs when the secondary mutation arises. At this point, say generation *t*, the individuals at generation *t* + 1 descend from an individual without a mutation with probability *p*, from an individual with the initial driver mutation with probability (1 + *s*_1_)*p* and with the secondary mutation with probability (1 + *s*_2_)*p*. If *n*_1_ and *n*_2_ are the number of individuals with the initial and secondary mutations respectively, then *p* is derived from the condition:

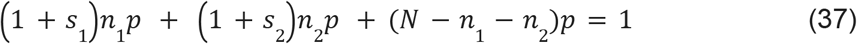

At generation *t*, the value of *n*_2_ is of course defined to be 1, but the procedure is iterated until the final time-point so the value of *n*_2_ can change.

This model can be easily extended to include a larger number of competing mutations, and is very similar to the one defined in the previous section that includes fluctuations in fitness values except in this case the selection parameter is inherited.

Assuming this model of multiple driver mutations, it is clear that rescaling the parameter values in the way described in this paper while keeping the fitness values of both clones fixed will result in trees that are indistinguishable. This is because scaling *L* or σ^2^ will naturally rescale the population size over time of both clones by the same factor, and the total population size that contains them *N* will also be scaled by the same factor. As a result, both clones will be found in similar proportions at the final time-point under any scaling of *c*, and any set of randomly sampled lineages will produce trees that are indistinguishable. Figure 10 demonstrates this idea using simulations (for details of simulations see next section).

**Figure 10.**
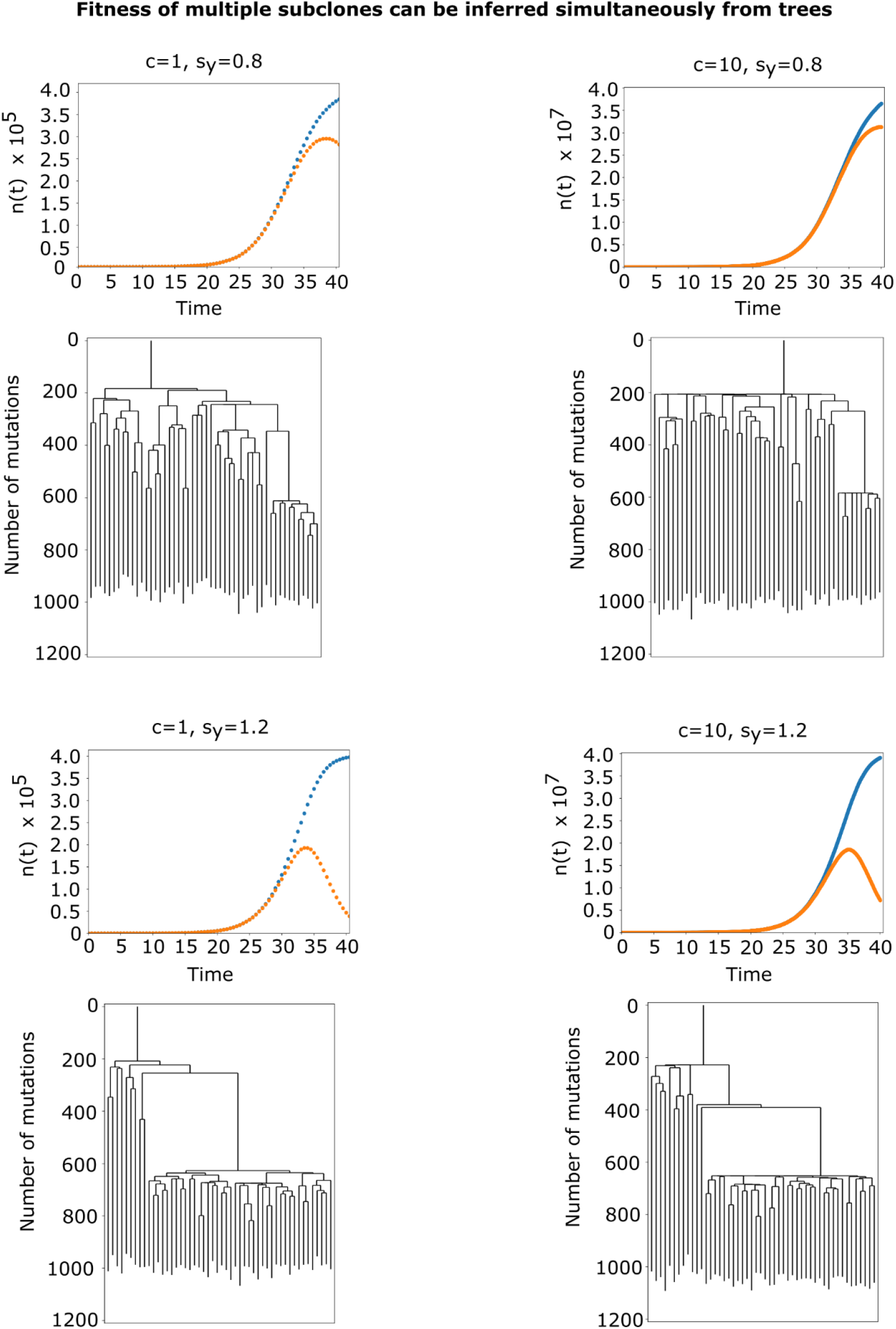
Trees constructed from populations with multiple competing mutations are similar even when the biological details are drastically different. Clonal expansions were simulated for 40 years and included a secondary driver mutation with increased fitness that occurred after 20 years of the initial expansion. Random bottlenecks at each generation were also imposed. Each of the 4 panels corresponds to different simulated expansions. For each of the four panels, the initial driver mutation had fitness *s*_*y*_ = 0. 4, the secondary driver mutation had fitness *s*_*y*_ = 0. 8 in the top two panels, but *s*_*y*_ = 1. 4 in the bottom two. The simulations corresponding to the right hand columns assumed 10 times as many generations and a bottleneck that was 10 times smaller than the left hand column so that the the population sizes over time of the subclones differed by a factor of 100 across both columns. The population sizes over time of each of the expansions were plotted in each panel, where the orange dots denote the number of mutants with the initial driver mutation, and the blue dots denote the number of mutants that have either the initial driver mutation or both mutations. For each expansion, 50 lineages were randomly sampled from individuals with either the first or secondary mutation and their trees are shown. As expected, the trees from the top row look qualitatively different from the bottom row since the fitness of the secondary mutation is different, but the trees from the left column are qualitatively similar to their corresponding trees on the right column because they share the same fitness values.

We can further extend this idea to scenarios where mutations that increase the fitness of a population are regularly introduced into a population. To do this, we can assume that selectively advantageous mutations are introduced into the population randomly per individual per generation with a mutation rate *U*. This mutation rate is different from the mutation rate earlier which refers to neutral mutations that do not provide a fitness advantage to the individuals in which they occur and are used only to construct the tree. Whenever a mutation arises in an individual, a value *q* is drawn from a distribution *D* that increases the selection value *s* of the individual that contains the mutation so that the new selection value is *s* + *q*. The *s* value of any given individual is therefore the sum of the *q* values accrued over time determined by the number of mutations accrued by its ancestral lineage and *D*. Under this model, we can also rescale the parameter values in a similar way as before, including the number of generations *L*, each of the selection values generated from *D* by a factor of 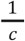, and the total population size *N* by a factor of *c*. Under this scaling, the fitness of each possible mutation introduced into the population in growth per year remains the same because of the increase in *L*, and although the decrease in *s* increases the probability of extinction by a factor of *c* of any mutation, the increase in *L* increases the rate at which the mutations are introduced into the population. As a result, the same number of mutations with equivalent fitness arise and establish in the population, but with population sizes over time scaled by a factor of *c*. Since *N* is also scaled by a factor of *c*, the individual subclones expand to similar proportions of the population and their rates of growth are identical to the unscaled process. As a result, we can infer the fitness values and times at which the different mutations arose from the lineage tree of a random sample of individuals at the final time-point without knowing the specific biological details, such as the number of generations, the distribution of number of offspring, or fluctuations in the timing between generations.

## Simulations

### Simulated trees show cancer and virus trees look similar whenever their fitness values are the same

In the previous sections, we have shown that fitness can be reliably inferred from lineage trees independent of the total number of generations and the value of σ^2^. If this is true, then a tree constructed from a cancer and from a virus should look identical whenever their fitness values are the same, but look different whenever their fitness values differ.

To show this, simulated example trees were constructed using realistic parameter values from MPN bone marrow cancers (parameter values obtained from [1], [2], [18]), and HIV replication (parameter values obtained from [6], [19], [20]), under different fitness values. In the simulated cancer expansions, the driver mutation was acquired at the 10^*th*^ generation, and the subsequent clonal expansion was simulated for an additional 25 generations. 1 generation (or equivalently one cell division) per year on average was assumed and the time between generations for each cell throughout the history of the expansion was drawn from a Gamma distribution with shape parameter = 5 and scale = 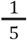. For each draw from the Gamma distribution, the cell would make a decision and either divide or undergo cell death with equal probability if it was wild-type, and divide with probability 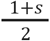 and undergo cell death with probability 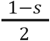 if it was a mutant. An average of 20 neutral, somatic mutations drawn from a Poisson distribution was inherited by each cell per cell division. The lineage tree was then constructed by randomly sampling 22 cancer cells at the final time-point. Two cancer trees were simulated using this procedure, one for *s* = 0. 1, and another for *s* = 0. 7. HIV growth dynamics were also simulated where cells are infected and sequential bottlenecks are applied. Similar to the cancer simulations, a mutation is acquired at generation 10 (the 10^*th*^replication cycle) by one of the viral particles and a subsequent clonal expansion is simulated for 25 generations. In contrast to the cancer 10 simulations, 1 generation is instead assumed to be 48 hours on average and is drawn from a Gamma distributed with shape parameter = 5. At the end of each generation, the HIV particle has a 10% chance of producing offspring. If it produces offspring it will produce 10^3^ offspring on 10^3^ average drawn from a poisson distribution if it is wild-type and 10^3^(1 + *s*) if it is a mutant. A bottleneck is then applied to the remaining progeny *X* by drawing the final number of progeny from ∼ *Bin*(*X*, 0. 01). Each viral particle inherits 30 neutral mutations on average drawn from a poisson distribution per replication event [20]. The lineage tree is then constructed by randomly sampling 22 viral particles at the final time-point. Two HIV trees were simulated using this procedure, one for *s* = 0. 1, and another for *s* = 0. 7. All four trees are plotted in Figure 1. As expected, the trees from both the cancer and virus look the same when their fitness is the same.

### Centered moments of simulated expansions match their theoretical values

To verify our derived expressions for the moments of the population size in the early drift phase, we show computationally that the k-th centered moment of the population size over time grow as 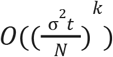. To do this, we simulated 10,000 clonal expansions according to the Wright-Fisher process with selection. The trajectories were each simulated for *L* = 200 generations and conditioned on surviving extinction at the final time point, and the parameter values used were *s* = 0. 1, *N* = 2^15^, and 1 generation per year was assumed. We then simulated another 10,000 trajectories identically to the first set of trajectories except we assumed 10 times as many generations so that 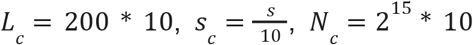 and assumed 10 generations per year. We then simulated a third set of 10, 000 trajectories in the same way as the first set of trajectories, but instead rescaled σ^2^ by a factor 10. This was accomplished by subsampling 10% of the total population size *N* at each generation using a hypergeometric distribution, and by letting the next generation of *N* individuals descend from the remaining wild type individuals that survived the bottleneck with probability *p*, and from the remaining mutant individuals that survived the bottleneck with probability (1 + *s*)*p*. The mean population size over time both with and without the appropriate offsets predicted to align the means and the *k* ^*th*^ central moments are plotted in Figure 6.

### The impact on the inferred age of onset when ABC model has incorrect assumptions

To verify our theoretical calculations that the impact on the inferred time of onset is at most one generation, we simulated 10 data trees and carried out ABC on them. Out of those data trees, we selected the tree where ABC most accurately inferred the true parameter values. The parameter values of that data tree were *g* = 50, *L* = 70, *s* = 0. 264911, *N* = 10^9^, *a* = 35 years, and the mutation rate was 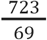. We then carried out ABC many times on the same data tree using *cL* generations for different values of *c*. The inferred age of onset was then plotted against *c* in Figure 7 to show that the offset in the inferred age of onset approaches 1 generation which is consistent with the theoretical bound derived above.

### Trees constructed from simulated expansions with multiple competing driver mutations look identical whenever the fitness values of each subclone are the same

Here we show that the coalescent structure of lineage trees only depends on the fitness values and time of occurrence of mutations even when multiple driver mutations are present. To do so, we simulated clonal expansions according to the Wright-Fisher process with selection, except that a secondary driver mutation occured within the expanding clone with a higher fitness value. In addition, we included bottlenecks at each generation similar to the bottleneck applied in the simulations in the previous section. We first simulated 5 clonal expansions where the initial driver mutations had fitness *s*_*y*_ = 0. 4, the secondary driver mutation had fitness *s*_*y*_ = 0. 8, and the first driver mutation expanded for 40 years while the secondary mutation expanded for the last 20. The duration of each generation was half a year. 20 neutral mutations were accrued per lineage per year, and a bottleneck of 50% out of a total population size *N* = 4 * 10^5^ was used. 50 lineages were randomly sampled at the final time-point from the set of individuals carrying either the first or secondary mutation and trees were constructed. We then simulated another 5 trees in the exact same way except in this case the secondary mutation instead had fitness *s*_*y*_ = 1. 2. Lastly, we simulated two more sets of trees, each set containing 5 trees as well, identically to the first two except that the number of generations was 10 times as large and the bottleneck size was 10 times as small. Therefore, the total population size *N* was 100 times larger because the population size of both clones is naturally rescaled by a factor of 100. From each set of trees, we chose 1 representative example and show them in Figure 10. The trees constructed from the populations having different fitness values for the secondary mutation look different, while trees constructed from the populations having similar fitness values for the secondary mutation look similar even though the number of generations and bottleneck sizes differ by a factor of 10 and the total population sizes over time of each subclone by a factor of 100. Lastly, here we have only chosen a representative example from each ensemble of trees. Although the structure of trees fluctuate within each ensemble, the ensemble of trees were qualitatively different when the fitness of the secondary mutation differed, but qualitatively similar when the fitness was the same.

### Inference of fitness and time of onset using simulated trees

In this section, we will use simulations to validate the above theoretical results on robustness of fitness inference to the assumptions in the model used for the inference. We will first briefly discuss a computational method for inference, and then show that we can reliably infer fitness and age of onset from simulated trees even when the biological details of the clonal expansions are drastically different from those assumed in the model for inference.

To infer fitness and age of onset from the trees of simulated clonal expansions, we use an algorithm called Approximate Bayesian Computation (ABC). To carry out ABC, we generate simulated trees under many different parameter values drawn from a prior distribution. The simulated trees are then compared to the data tree using a metric distance. If the distance between a simulated tree and the data tree is smaller than a predefined threshold, the parameter values associated with the simulated tree are kept, and otherwise discarded. After many iterations, we obtain a posterior distribution of parameter values that capture the best estimates of the ground truth parameter values in the data tree.

In our particular implementation of ABC, we use the Wright-Fisher model with selection to generate clonal expansions that arise when a mutation is acquired by an individual in the population randomly at some generation. Neutral mutations are also accrued along the lineages by drawing the number of mutations from a Poisson distribution with a mean of ~20 mutations per generation. 22 cells are randomly sampled at the final time-point, and the lineages are traced backwards in time to construct a tree where the branch lengths are measured in number of mutations. The simulated tree is then compared to the data tree using a metric distance. To compute the distance, we first convert each tree to an LTT curve, which plots the number of extant lineages as a function of time expressed in number of mutations. The distance between two lineage trees corresponds to the area between their LTT curves.

To verify our theoretical results, we generated data trees by simulating clonal expansions where s was drawn independently for each mutated individual from a uniform distribution on (0, 2 * *s*). In addition, 10 percent of the individuals at each generation were killed so that the individuals in the next generation descended from only 90 percent of the population increasing the variance in the number of offspring per individual. In contrast, the model used to generate the simulated trees to perform ABC assumed a constant value of s for each individual and did not kill any fraction of individuals. In addition, the model used for the inference incorrectly assumed that the number of generations were 10 times smaller than the number of generations in the data. Figure 11 shows the inferred value of fitness, the times in years at which the mutations were acquired, and the number of mutant individuals at the final time-point.

**Figure 11.**
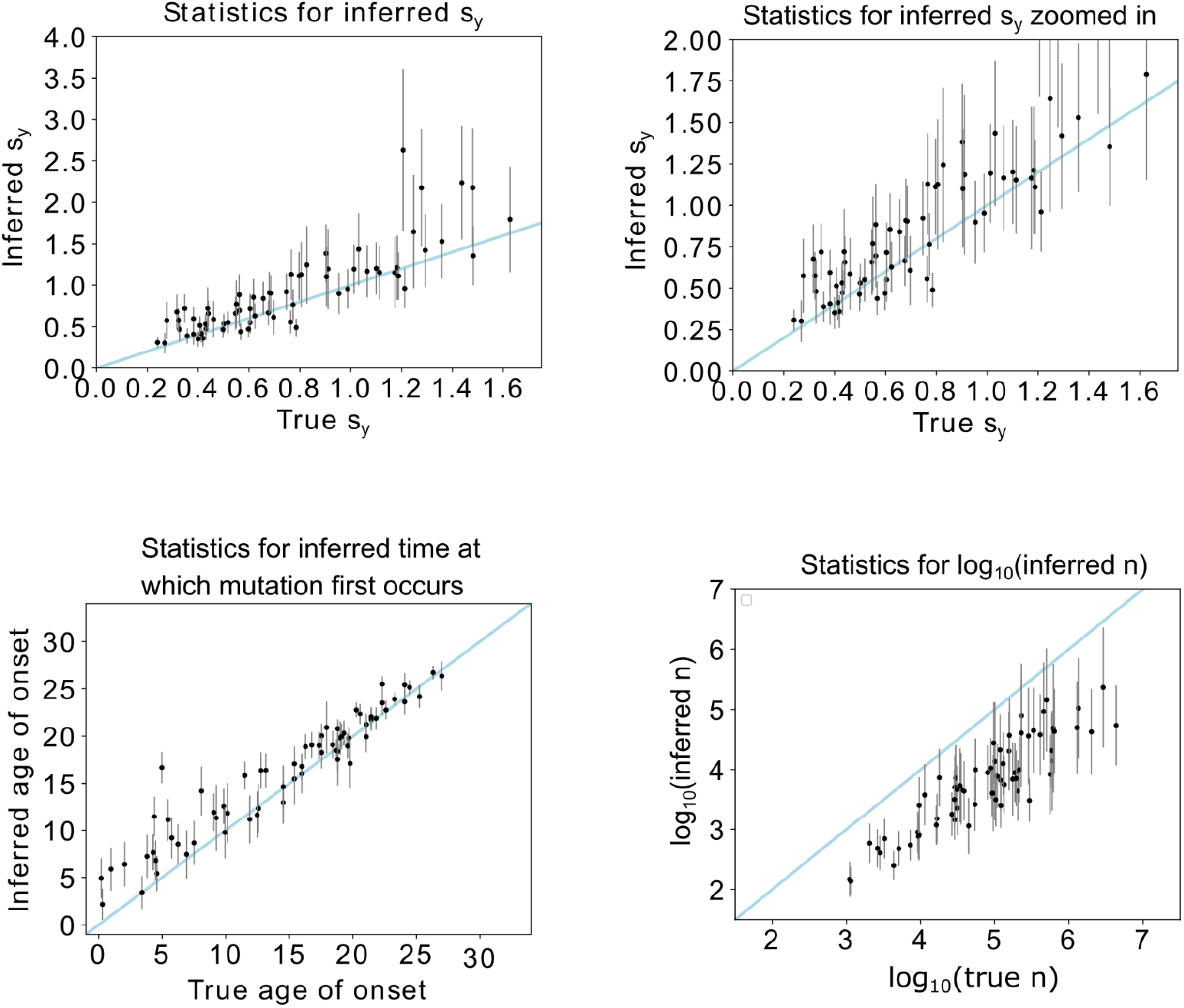
Inferences from simulated data trees where the biological details of the model used for the inference were completely different from the model used to simulate the data trees. Many data trees were simulated under different parameter values and ABC carried out on each one to infer the fitness, age of onset, and final population size. Data trees were generated using 10 times as many generations as assumed in the ABC, which was *L* = 35. The generations occur over a total time of *a* = 34 years. Heterogeneity was included in the selection parameter by drawing its value from a uniform distribution for each individual of the clone, but the average fitness was kept a constant value. Furthermore, at each time step, 10% of the population was randomly killed so that each generation descended from 90% of the individuals in the previous generation increasing the variance in the number of children σ^2^. The ABC model did not include this fitness heterogeneity and increased variance. The average fitness, number of generations, and time of onset was drawn randomly and the corresponding value used to simulate each data tree. Inferences were carried out with the ABC model that did not incorporate this heterogeneity and the values were plotted. Each plot shows the means of the inferred vs true parameter values for all the inferences shown as dots with error bars denoting 1 std. As expected, fitness and age of onset can be reliably inferred, while the number of mutant individuals at the final time point (show in the bottom right panel) are under inferred since the expansions in the data fluctuated to a larger population size because of the increased number of generations and a larger σ^2^.

As expected by our theoretical calculations, we are able to correctly infer the fitness and the time at which the mutation arose despite the incorrect assumption in the models used for the inference. Also as expected, the inferred number of mutant individuals at the final time-point is smaller than the true value. This makes sense because the true model had a smaller average growth rate per generation s and a larger σ^2^ compared with the inference model. Therefore, the population in the true model experienced larger fluctuations and had to drift to a larger size before exponentially growing. Importantly, despite the fact that the inferred population size is a scaled version of the true value, the inferred fitness matches the true value.

## Discussion

In this paper we have shown that fitness and age of onset can generally be inferred from the lineage trees of a random sample at the final time-point from a population that has grown exponentially, independent of the underlying biological details. The important limitation is that the population must be clonal or consist of multiple expanding clones where the stochastic behavior of each mutant individual at each generation within each expanding clone is identical. This assumption could break down if the σ^2^ value or the timing between generations changed rapidly over time within an individual clone so that individuals in the later history after escaping extinction had very different statistics. This assumption can also be broken if selectively advantageous mutations are introduced extremely rapidly into the population so that new mutations arise within any given clone before the clone has the opportunity to establish and escape extinction. This is because our results rest on the assumption that a clone escapes extinction after reaching population size 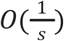 which could break down considerably if it accrued many selectively advantageous mutations beforehand. We have also not considered mathematically the impact on trees when the timing between generations takes on the form of an atypical distribution, such as a power-law which could conceivably model dormant cancer cells for example.

A further limitation of this work is that we have ignored the noise introduced by fluctuations in the number of mutations per generation per individual when constructing trees. We have also assumed that the mutation rate is constant, or at the very least that we have measurements of the mutation rate as a function of time which can be used to convert the branch lengths of a tree from number of mutations to absolute time. Lastly, we are assuming that whatever model we are using can correctly account for the biological scenarios we are trying to infer. For example, if there are multiple driver mutations within a cancer expansion and the model used for the inference only incorporates a single driver mutation, then the model clearly cannot account for the fitness value of the multiple mutations. Importantly, our results do not tell us what biological scenarios to model, but merely that once we have chosen the biological scenario to model, we don’t need to get the details of replication correct.

In this work, we also made some explicit mathematical assumptions which we will further highlight. In particular, the Fokker-Planck and stochastic differential equation approximation to the Wright-Fisher process are derived in a large *N* and small s limit. Furthermore, the expressions for deriving coalescent statistics are also derived in a large population size limit, and so there may be some deviations to the derived tree statistics that occur in the initial phase of the clonal expansion when the number of mutant individuals is small. Minor deviations of this sort were observed by eye in some of the trees but likely will not have a large impact on the inferred values as seen in Figures 5 and 8. However, any large deviations from these limits will impact the statistics of trees.

Even with these caveats in mind, it appears that fitness and the timing of mutations can generally be reliably inferred from lineage trees, even with little knowledge of the biological details and a large amount of heterogeneity included. As a result, using lineage trees to infer fitness and the timing of mutations in exponentially growing clones is likely robust and a reliable method for inference.

## Acknowledgements

This work was supported by the National Institutes of Health (NIH) grants NHLBI R01HL158269 and NIH T32GM135014. We also acknowledge funding from AACR-MPM Oncology Charitable Foundation Transformative Cancer Research Grant, Gabrielle’s Angel Foundation for Cancer Research, and the V Foundation for Cancer Research.

